# Coordinated acetylcholine release and adaptation of neuronal representations in the retrosplenial cortex during contextual uncertainty

**DOI:** 10.64898/2026.05.02.722331

**Authors:** Daniel Goodwin, Ani Boja, Charline Tessereau, John B. Issa, Guochuan Li, Yulong Li, Daniel Dombeck, Michael Rule, Jonathan T. Brown, Jack R. Mellor, Jonathan Witton

**Affiliations:** Department of Clinical and Biomedical Sciences, Faculty of Health and Life Sciences, University of Exeter, UK; School of Physiology, Pharmacology and Neuroscience, University of Bristol, UK; School of Engineering, Mathematics and Technology, University of Bristol, UK; Champalimaud Foundation, Lisbon, Portugal; Max Plank Institute for Biological Cybernetics, Tuebingen, Germany; Department of Neurobiology, Northwestern University, Evanston, Illinois, USA; State Key Laboratory of Membrane Biology, School of Life Sciences, Peking University, Beijing 100871, China; Peking-Tsinghua Center for Life Sciences, New Cornerstone Science Laboratory, Academy for Advanced Interdisciplinary Studies, Peking University, Beijing 100871, China

**Keywords:** Acetylcholine, position cells, reward cells, uncertainty, retrosplenial cortex

## Abstract

Accurate learning underpins optimal predictions and requires the brain to navigate uncertainty, identifying important information and distinguishing between expected and unexpected uncertainties. Theoretical models propose that the neuromodulator acetylcholine positively correlates with expected uncertainty, priming neuronal networks for learning. We tested this hypothesis by measuring acetylcholine release and neuronal representations in the retrosplenial cortex of mice whilst challenging them with expected and unexpected uncertainties of reward location. Acetylcholine release did not directly correlate with expected or unexpected uncertainty but instead increased with changes in expected uncertainty. In tandem, increased expected uncertainty shifted neuronal representations from a positional reference frame to focus on salient landmarks, such as reward location. Transitions in expected uncertainty also accelerated remapping of neuronal representations and behavioural adaptation when unexpected uncertainty was experienced. Thus, we demonstrate acetylcholine release in retrosplenial cortex discerns types of uncertainty and correlates with learning speed in uncertain environments.

## Introduction

Plasticity is a fundamental and vital feature of brain function enabling learning and behavioural adaptation. New information needs to be rapidly incorporated into existing estimates and predictions that are encoded within neuronal networks, but in real world situations this updating depends on uncertainty. For example, if we find our favourite ice cream van parked on the same road every day there can be some variation in the exact location, but this is *expected uncertainty* and, once we have learnt the range of variation, does not require any great adjustment of predictions to find our ice cream. However, if one day the ice cream van has parked at the far end of the street or on the next block, this is outside the boundaries of our predictions and is therefore *unexpected uncertainty,* requiring an update in estimates of the van’s future location and adjustment of behavior, such as budgeting extra time to find the van or selecting a more reliable vendor. Importantly, there is a dynamic interaction between expected and unexpected uncertainties, since the degree of expected uncertainty dictates the threshold for detecting unexpected uncertainty, and conversely, an unexpected event will increase the level of expected uncertainty ^1,2^.

To distinguish between these forms of uncertainty and engage accurate learning the brain must provide signals that link the uncertainty condition to learning mechanisms. Longstanding evidence shows that neuromodulator systems provide signals associated with expectation, error, and uncertainty, with their release reconfiguring neuronal circuits to enable the update of estimates and memories ^3,4^. This suggests neuromodulators determine the importance and impact of new information in a manner that incorporates estimates of uncertainty. Acetylcholine has a particular prominence in this regard since its release promotes plasticity and learning ^5,6^ and is associated with arousal, attention, and expected uncertainty ^7,8^. Along with other neuromodulators such as noradrenaline and dopamine, acetylcholine is also associated with surprise or unexpected uncertainty that necessitates a shift in inference about the current environmental state ^9–12^.

Changes in expected uncertainty are predicted to alter the environmental context, as knowledge about stimulus-outcome contingencies becomes unreliable for guiding effective behaviour. This process is proposed to be accompanied by acetylcholine release, which facilitates plasticity to support learning of revised contingencies ^1,4^. Changes in expected uncertainty can arise when the reliability of known environmental features shift within an otherwise stable context. However, unexpected uncertainty also necessitates learning of revised contingencies and therefore may also require signals to facilitate plasticity. Furthermore, because expected and unexpected uncertainties interact, unexpected events that alter stimulus-outcome contingencies may also elevate expected uncertainty as new contingencies are learned. Notably, despite these predictions, two central assumptions of expected uncertainty remain largely untested: firstly, that acetylcholine release tracks expected uncertainty rather than unexpected uncertainty, and secondly, that changes in levels of expected uncertainty require plasticity to update neuronal representations of environmental context.

To test these hypotheses, we measured acetylcholine release and neuronal representations during conditions of expected and unexpected uncertainty in the dysgranular retrosplenial cortex (dRSC) of mice, a region important for linking contextual information with behavioral decisions ^13,14^. The dRSC receives dense cholinergic innervation from the medial septum/diagonal band nuclei and nucleus basalis of the basal forebrain ^15–18^ and houses representations of salient environmental features, including spatial position ^19–21^, proximity to boundaries ^22–24^ and visual landmarks ^25–27^, and goal-related variables such as reward locations^24,27,28^ and reward value ^29^.

While egocentric (self-referenced) components of dRSC representations are context invariant ^23^, allocentric (externally referenced) positional and reward representations dynamically adapt between contexts ^14,29–31^. In novel environments, these representations undergo global updating, similar to the remapping of place cell activity in the hippocampus ^32^. Indeed, the formation of dRSC positional representations depends on hippocampal input ^26,33^, but once established, these representations can be reactivated independently of the hippocampus ^34,35^. Additionally, the RSC supports transformations between egocentric and allocentric frames of reference ^36^ making it an ideal brain structure to investigate interplay between intrinsic predictions of uncertainty and environmental shifts.

We found that acetylcholine release increases with transitions in the level of expected uncertainty related to reward location. In contrast, acetylcholine is only released during unexpected uncertainty when it is associated with a contextual change that also alters expected uncertainty, such as exposure to a novel environment or during a transition from high to low expected uncertainty. Consistent with predictions, neuronal representations did adapt substantially to increased expected uncertainty, with a recruitment of cells to represent the uncertain reward location, but not other spatially stable features of the environment. Furthermore, neuronal representations and behaviour adapted faster to unexpected uncertainty when they were accompanied by a transition from high to low expected uncertainty that triggered acetylcholine release. Thus, our data identify the dRSC as a locus of uncertainty processing and provide a resolution to longstanding questions regarding the role of acetylcholine coordinating plasticity in conditions of uncertainty.

## Results

### Expected and unexpected uncertainty of reward location

To investigate the role of acetylcholine in regulating plasticity of neuronal dRSC representations under different conditions of uncertainty, we first developed the Uncertain Reward Task (URTask) for head fixed mice running in a linear virtual reality environment ^37^. In each daily session, mice ran multiple traversals of a 3 m track with a reward delivered at a single location on each traversal. At the end of each traversal, the mice were teleported back to the start after a 3 s timeout which marked the end of the previous trial and the start of the next one. The URTask is designed to manipulate the degree of expected and unexpected uncertainty by changing the distribution of possible reward locations on the track. In the initial condition, the reward location was consistently in a narrowly defined location (either 90 cm or 100 cm along the track) – a condition of low uncertainty once mice were well trained. From this baseline, expected uncertainty was increased by increasing the variability of the reward location on a lap-by-lap basis. This was implemented by increasing the size of the zone in which reward was delivered in any individual track traversal, such that a single reward was delivered per traversal with equal probability at one of 10 equally spaced locations within the broader (100 cm wide) reward zone (Fig. 1A-C). In contrast, unexpected uncertainty was induced by moving the reward location permanently to a new location (Fig. 1D-F). Importantly, there is potential overlap in the induction of expected and unexpected uncertainties such that a) when expected uncertainty is increased by broadening the reward zone there may also be some unexpected uncertainty induced as the new reward location distribution is learnt (Fig. 1A) and b) when unexpected uncertainty is induced by permanently moving the narrowly defined reward zone this infers a change in the task state where such manipulations are now possible and therefore expected uncertainty increases (Fig. 1D). Furthermore, the level of expected uncertainty may alter the detection and response to unexpected uncertainty. This was tested in the URTask by moving the reward location from a broad reward zone with high variability to a new single permanent zone in a different position along the track (Fig. 1G-I), which is predicted to induce less unexpected uncertainty than when the initial reward zone is narrow and also to reduce expected uncertainty. We term this transition ‘uncertainty interaction’.

Behaviorally, mice showed consistent abilities to learn and adapt to changes in reward locations and we found no differences between male and female mice (Fig. S1). Although reward delivery was not contingent on behaviour, all trained mice demonstrated anticipatory licking and velocity deceleration on approach to the reward and mice then adapted to a new reward location after a switch (Fig. 1). After at least 10 days of training, mice ran at least 40 traversals of the track in each session.

We complemented the URTask with a switch to a novel visual environment – a commonly used manipulation that induces a large contextual shift inducing both unexpected uncertainty and increased expected uncertainty of potential future environmental change (Fig. 1J-L). This manipulation has been shown to cause complete remapping of spatial representations in dRSC^32^. Since the reward zone is moved at the novel environment transition, behavioral adaptation of mice to the novel environment was similar to moving the reward zone.

**Figure 1.**
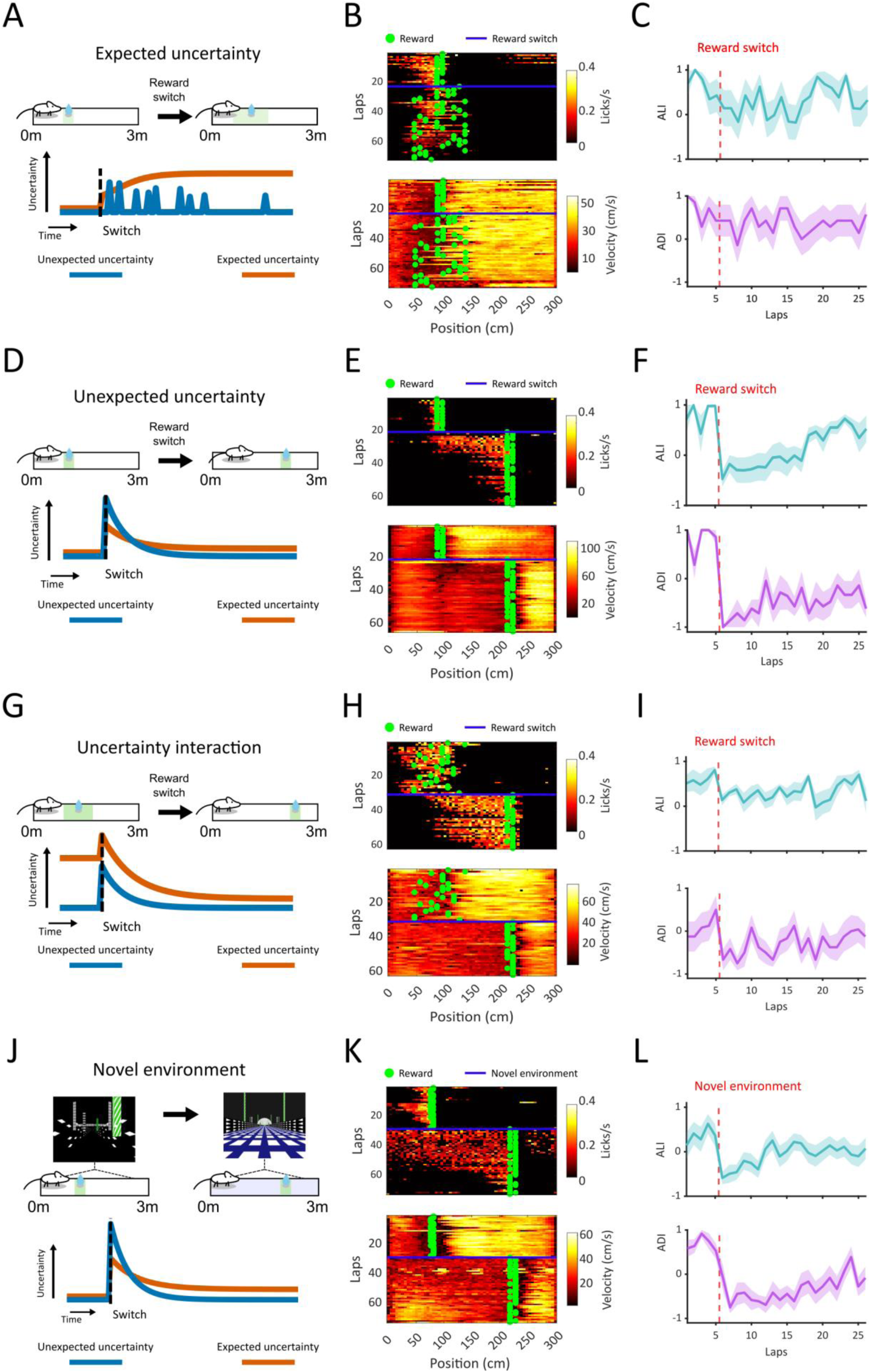
Behavioral adaptation to uncertainties in reward location in the Uncertain Reward Task (URTask). (A) Expected uncertainty is increased by changing the reward zone from a narrow predictable location to a broader less predictable location where reward can be delivered once anywhere within the zone on each lap. Visual cues remain unchanged throughout the session. Below are predicted levels of expected and unexpected uncertainties: a switch to increased expected uncertainty coincides with small, transient increases in unexpected uncertainty as the mouse learns the new task state. (B) Example mouse behavior measured by lick frequency (top) and running velocity (bottom) indicating anticipatory licking and deceleration upon reward approach and adaptation to new reward location contingency. (C) Average anticipatory lick index (ALI) and anticipatory deceleration index (ADI) before and after introduction of expected uncertainty (reward switch, dashed line) (N=7 mice). (D-F) Same as A-C but for unexpected uncertainty induced by switching the reward location to a new consistent location (N=7 mice). (G-I) Same as A-C but for uncertainty interaction tested by starting the session in high expected uncertainty with a broad reward zone and moving to a new consistent reward location with resulting reduction in expected uncertainty (N=8 mice). (J-L) Same as A-C for teleport to a novel visual environment with a new reward location (N=12 mice).

### Acetylcholine release and neuronal plasticity in response to a novel environment

Acetylcholine release in the hippocampus increases when mice are exposed to a novel environment ^38^, while exposure to novel environments also causes remapping of neuronal representations in both the hippocampus and dRSC ^32,38^. Encountering a novel environment also constitutes a large contextual change that generates unexpected uncertainty and an increase in expected uncertainty. To test the relationship between acetylcholine release and neuronal plasticity in dRSC, we first measured acetylcholine release and recorded the activity of neuronal populations in dRSC in response to exposure to a novel environment.

We expressed the fluorescent acetylcholine sensor GRAB-ACh4l ^38,39^ or Ca^2+^ indicator jGCaMP8m ^38,40^ in dRSC neurons and monitored their dynamics using two-photon microscopy whilst mice ran laps of the virtual reality track (Fig. 2A-C).

GRAB-ACh4l fluorescence (ΔF/F) was positively modulated with running velocity and plateaued at higher velocities (Fig. 2D,E, Fig. S2), suggesting saturation of the sensor or acetylcholine concentration, consistent with previous findings ^38,39^. Fluorescence signals in response to velocity were abolished in control mice expressing the ligand-insensitive sensor variant GRAB-ACh3.0mut ^39^, confirming that GRAB-ACh4l reports acetylcholine release within dRSC (Fig. S2). However, in contrast to the velocity signal, we did not detect any GRAB-ACh4l fluorescence signal in response to reward (Fig. S2). To sensitively detect bidirectional changes in acetylcholine release while avoiding signal saturation at higher running velocities and accounting for velocity modulation of the signal, we analysed GRAB-ACh4l fluorescence filtered at 0-10 cm/s and calculated velocity-controlled averages (Fig. S2). We found velocity controlled GRAB-ACh4l fluorescence lap-averages were stable across control recording sessions where no manipulation of the environment occurred, demonstrating limited bleaching or changes in acetylcholine release across the duration of the recording session (Fig. S2).

Introducing both unexpected and increased expected uncertainty via teleportation to a novel virtual environment elicited a rapid increase in acetylcholine release over and above the velocity modulated acetylcholine release dynamics (see Methods). This novel environment induced increase remained elevated throughout the recording session (Fig. 2E-G; Table S1). Additionally, GRAB-ACh4l fluorescence dynamics were consistent across layers I and II/III of dRSC during baseline conditions in a familiar environment and following exposure to a novel environment, with no layer dependent signal for either velocity or novelty (Fig. S3), similar to observations in the hippocampus ^38^. These results show that acetylcholine release across multiple layers of dRSC is associated with distinct behavioral contexts: movement and novelty.

jGCaMP8m signals in dRSC neurons revealed positionally modulated neuronal activity where individual cells’ activity tiled the entire length of the track, as previously reported for dRSC populations in linear environments ^19,27,32,41^. Positional representations were stable over the duration of a recording session and consistently over-represented reward locations and the ends of the track (Fig. 2H; Fig. S4). Switching the visual environment from a familiar to a novel track caused positionally correlated cells (PCCs) to globally remap, and a new representation of the novel environment emerged which retained some representation of the ends of the track (Fig. 2H-J). Similar to the hippocampus ^37,42–44^, a significant minority of cells moved their peak activity location in line with the reward location change, quantified by aligning activity on each lap to the precise reward location. These reward correlated cells (RCCs) were selectively active within a zone ± 25cm of reward (6.7 ± 2.4% of cells, Fig. 2K,L) and were not detectable at other regions of the track (8.1 ± 0.4% of cells on the reward reference frame away from the reward zone, Fig. 2K,L) indicating that these cells were primarily responsive to the reward location rather than animal position or visual features of the environment.

Together, these findings demonstrate that experience of a novel environment drives a rapid and sustained increase in acetylcholine release within the dRSC, accompanied by substantial remapping of the neuronal representation of the environment. This pattern is consistent with a role for acetylcholine in supporting experience-dependent plasticity in the dRSC. However, neuronal representations in a familiar environment remained stable despite coinciding with acetylcholine release associated with movement.

**Figure 2.**
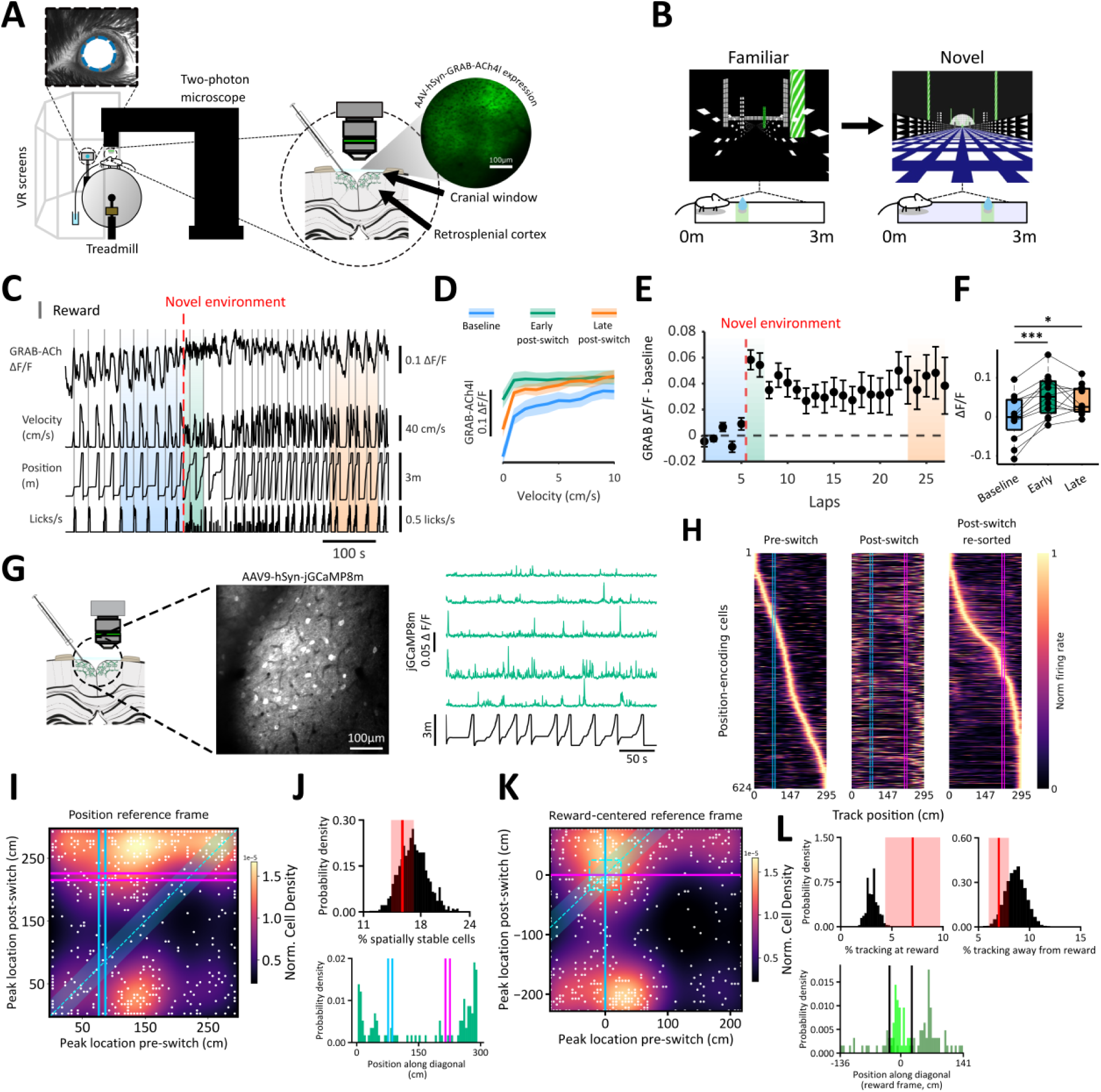
Increase in acetylcholine release and neuronal representation plasticity in response to a novel environment. (A) Experimental setup. Head-fixed mice navigated a linear virtual reality (VR) track projected onto screens surrounding a cylindrical treadmill. Forward and reverse treadmill movement controlled virtual position. Pupil dynamics were monitored using an infrared camera (inset; example pupil outlined in blue). The right schematic shows the cranial window over dysgranular retrosplenial cortex (dRSC) and an example two-photon field of view expressing GRAB-ACh4l. (B) Novel environment paradigm. Mice initially navigated a familiar virtual environment (left) and were then teleported to a novel environment (right) during the session. (C) Representative recording during an environment switch showing GRAB-ACh4l fluorescence (ΔF/F), running velocity (cm/s), track position (m), and licking rate (licks/s). Grey vertical lines denote reward delivery; red dashed line indicates the switch to a novel environment. (D) Relationship between acetylcholine signal ΔF/F) and running velocity (0–10 cm/s) across behavioural epochs: baseline (blue), early post-switch (green), and late post-switch (orange). N = 12. (E) Lap-by-lap baseline-subtracted ΔF/F aligned to the novel environment onset (red dashed line). Shaded regions indicate analysis epochs: baseline (blue), early post-switch (green), and late post-switch (orange). Black dashed line denotes ΔF/F = 0. (F) GRAB-ACh4l ΔF/F increases in a novel environment at early and late post-switch epochs. Repeated-measures ANOVA with Greenhouse–Geisser correction, followed by Dunnett-adjusted post hoc comparisons against baseline (\*\*\**p* < 0.001, \**p*<0.05). (G) Calcium activity was recorded from jGCaMP8m-expressing neurons. Example imaging field of view (left), with example ΔF/F traces from five neurons, aligned to VR position (right). (H) Normalized spatial activity maps for PCCs in the dRSC across the novel environment switch. Left, pre-switch activity maps sorted by peak firing location (reward location indicated by blue lines). Middle, post-switch activity maps for the same neurons sorted by pre-switch peak location (previous reward depicted by dashed blue lines, new reward by magenta lines). Right, post-switch activity maps reordered by peak firing position. (I) Galaxy plot illustrating the relationship between pre- and post-switch peak firing locations for individual neurons (white dots). Kernel density estimate (KDE) heat map indicates population density. Pre- and post-switch reward locations are indicated (blue and magenta lines, respectively). The cyan dashed diagonal and shaded band represent a ±25 cm stability threshold around unity; neurons within this band are classified as spatially stable. (J) Spatial shuffle analysis of stability across the environment switch. Top, null distribution, with the observed proportion of spatially stable cells indicated in red. This was not significantly different from the shuffled distribution (*p* = 0.78; one-sided permutation test). Bottom, probability density of neurons along the diagonal axis of the galaxy plot in (I), with pre- and post-switch reward zones indicated (cyan and magenta, respectively). (K) Reward-referenced galaxy plot for the environment switch. Peak firing locations are expressed relative to the reward location (zeroed per trial). White dots show individual neurons and KDE shading indicates density. RCCs are defined as peaks occurring within ±25 cm of reward.. (L) Distribution of reward-tracking neurons along a reward-referenced coordinate frame. Top and middle, probability density of neurons tracking reward location (“at reward”) or displaced from reward (“away from reward”). Bottom, distribution of reward-correlated neurons projected along the diagonal axis of a radial reward reference frame; vertical black lines denote ±25cm around reward. Reward tracking at the reward location was significantly above chance (*p* = 0.0002), whereas tracking away from reward was not (*p* = 0.9434; one-sided permutation tests).

### Acetylcholine release and neuronal plasticity in response to expected uncertainty

Having established the sensitivity of acetylcholine release and neuronal representations in dRSC to unexpected and high expected uncertainty elicited by a novel environment, we next tested cholinergic and neuronal population responses to specifically altering the level of expected uncertainty of reward location in the URTask (Fig. 1A-C). Increasing expected uncertainty is predicted to increase acetylcholine release and promote learning of the altered contextual state (e.g., cue reliability), but not to otherwise change neuronal representations of features of the context that remain stable ^1,7^. In a familiar environment, mice were first trained under low uncertainty, with a single predictable reward delivered within a 10 cm wide zone. In the test session, mice completed at least 20 laps in the low uncertainty condition before the reward location contingency was abruptly changed so that the single reward on any given lap was delivered within a 1 m wide zone, still centred at the same position along the track (Fig. 3A). Thus, the only manipulation was a decrease in the predictability of reward location, experienced as an increase in expected uncertainty once the new reward location contingency was learnt.

Because the manipulation increased the reward location distribution without shifting the average reward location, we observed limited behavioral adaptation. Mice exhibited a small decrease in anticipatory licking, with anticipatory deceleration remaining consistent (Fig. 1C, 3B). Due to the random variability of the reward location after the reward contingency switch, the mice did not always detect the manipulation immediately. However, within several laps, mice reliably exhibited an increase in pupil area that further increased over successive laps, indicating robust detection of the contingency change and a progressive increase in arousal that was independent of running velocity (Fig. 3C-E). This gradual increase in arousal mirrors the predicted increase in expected uncertainty as the more variable reward location is learnt (Fig. 1A).

Acetylcholine release measured by GRAB-ACh4l fluorescence also gradually increased across laps, mirroring the rate of pupil dilation and predicted levels of expected uncertainty (Fig. 3C-E; Tables S1 & S2). Interestingly, alignment of GRAB-ACh4l fluorescence to onset of movement or reward delivery revealed that whilst velocity, pupil dilation, and acetylcholine release were all positively correlated and coordinated at the onset of movement, reward delivery resulted in pupil dilation and reduced velocity with a corresponding decrease in acetylcholine release (Fig. S2). This dissociation indicates that the observed increase in acetylcholine release after reward contingency manipulation reflects increased expected uncertainty, rather than reward delivery per se.

**Figure 3.**
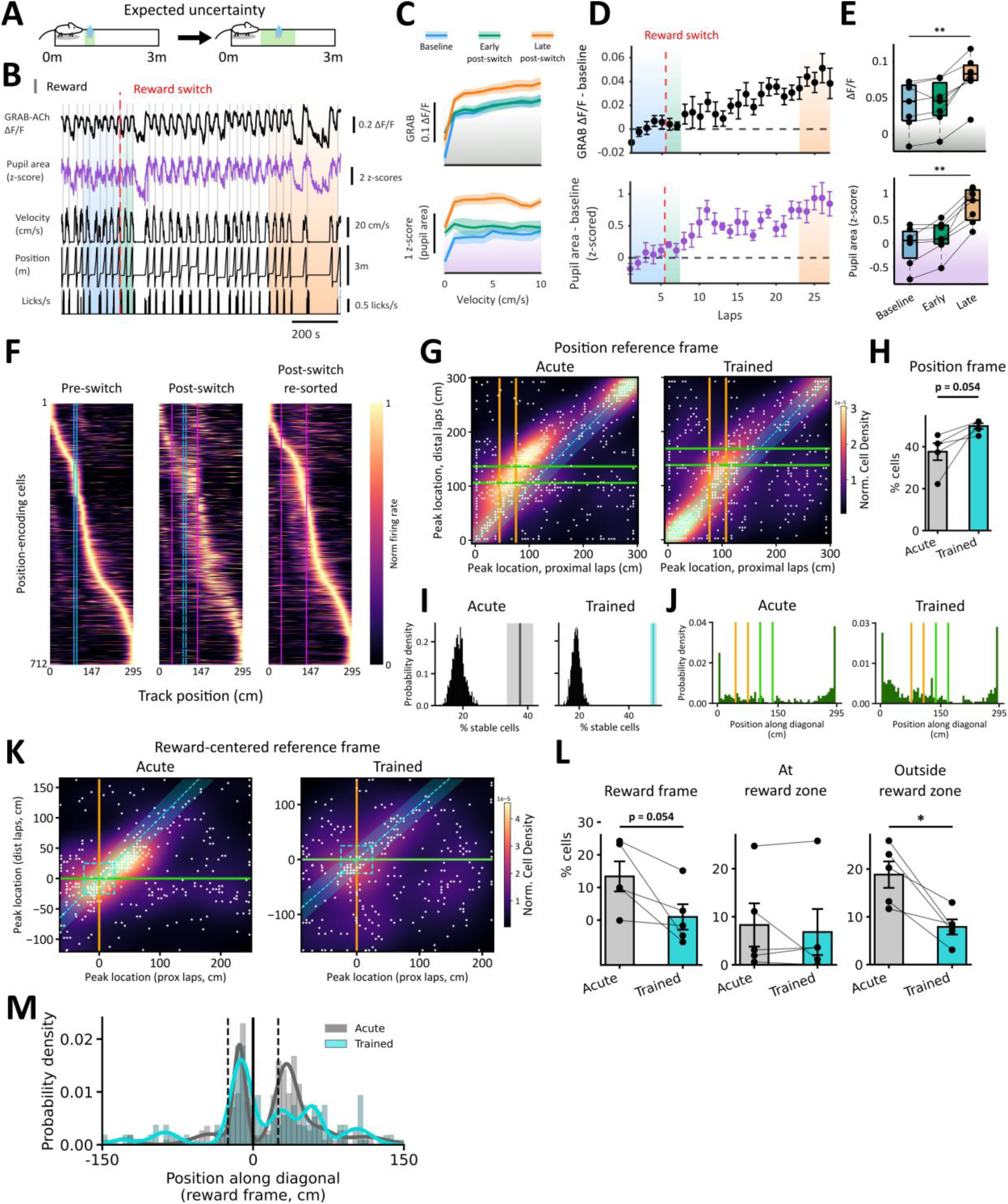
Acetylcholine release and neuronal representations in response to an increase in expected uncertainty. (A) Reward switch in which a narrow (10 cm) reward zone transitioned into a broad (1 m) reward zone. (B) Representative recording showing GRAB-ACh4l fluorescence (ΔF/F), pupil area (z-scored), running velocity (cm/s), track position (m), and licking rate (licks/s). Red dashed line indicates the reward switch. Transient increases in pupil diameter at the end of each lap coincide with temporary blackout of the visual display during the 3 s inter-trial interval. (C) ÄF/F (top) and pupil area (bottom) binned by running velocity (1 cm/s bins). Separate lines indicate baseline (blue), early post-switch (green), and late post-switch (orange) epochs. N = 7. (D) Lap-by-lap baseline-subtracted ΔF/F (top) and pupil area (bottom) aligned to the reward switch (red dashed line). Shaded regions denote baseline (blue), early post-switch (green), and late post-switch (orange) epochs. Dashed lines indicate zero. (E) Mean ΔF/F (top) and pupil area (bottom) across behavioural epochs (baseline, early, and late). Repeated-measures ANOVA with Greenhouse–Geisser correction, followed by Dunnett-adjusted post hoc comparisons against baseline (\*\**p* < 0.01). (F) Normalized spatial activity maps for PCCs. Left, pre-switch activity maps sorted by peak firing location. Middle, post-switch maps for the same neurons aligned to pre-switch ordering. Right, post-switch maps reordered by peak location. Cyan lines indicate pre-switch reward location, magenta lines denote post-switch span. (G) Galaxy plots showing the relationship between peak firing locations in proximal (orange) versus distal (green) laps (first vs. last third of reward zone post-switch). Left, acute condition (immediately following switch); right, trained condition (after 3 sessions with expected uncertainty). White dots represent individual neurons; colour indicates population density (KDE). Cyan dashed diagonal and shaded band denote ±25 cm stability threshold. (H) Percentage of spatially stable cells in acute versus trained sessions (paired t-test). (I) Spatial shuffle analysis corresponding to (G). Null distributions for acute (grey) and trained (blue) conditions. Observed stability was significantly greater than chance for both sessions (*p* = 0.001; one-sided permutation tests). (J) Probability densities along the diagonal axis for acute (grey) and trained (blue) conditions. Proximal reward location shown as yellow lines, distal as green lines (K) Reward-referenced galaxy plots comparing proximal (orange) and distal (green) laps in acute (left) and trained (right) conditions. Peak firing locations are expressed relative to reward position. Spatially stable cells are excluded. Cyan dashed diagonal indicates reward-aligned tracking. (L) Proportion of RCCs in acute and trained conditions. Left, total tracking; middle, tracking at the reward location; right, tracking away from reward (paired t-tests; *P < 0.05). (M) Distribution of neurons along the diagonal axis in the radial reward-referenced coordinate frame for acute (grey) and trained (blue) conditions. Gaussian mixture models fitted to each distribution. Black line indicates reward location; dashed lines denote ±25 cm around reward.

We next investigated how an increase in expected uncertainty affected neuronal representations in dRSC and how this evolved with experience. Unlike exposure to a novel environment, increasing expected uncertainty by increasing the reward location distribution did not induce global remapping of dRSC neuronal representations (Fig. 3F). To quantify the extent of remapping and assess the contribution of PCC and RCCs, we compared cellular activity on laps where the reward was delivered in the initial third of the reward zone (‘proximal laps’) versus laps where the reward was delivered in the final third (‘distal laps’). This separated PCCs that stably represented track position and RCCs that moved with reward location. 42.6 ± 5.4% of cells were PCCs after the initial increase in expected uncertainty and this increased slightly to 57.3 ± 2.1% in animals that were trained for at least 3 days on a track where the reward zone was still 1 m wide but 75 cm – 175 cm along the track (Fig. 3G,H). In both cases, the PCCs were concentrated at the ends of the track and the region leading up to the reward zone, with little representation between the reward zone and the end of the track (Fig. 3G). To assess RCCs, we again aligned cell activity to the reward delivery location for each lap. This revealed a high number of RCCs when expected uncertainty was initially increased (29.4 ± 5.4%) which then reduced after the mice were trained in the high expected uncertainty condition (19.9 ± 5.2%) (Fig. 3I-K). This training-related reduction was mostly accounted for by a reduction in RCCs that were active within the reward reference frame but not close to the reward location (>25 cm; ‘away from reward’ RCCs), whereas the percentage of RCCs active close to the reward location (≤25 cm; ‘at reward’ RCCs) remained consistent (Fig. 3L). Thus, there is plasticity of dRSC neuronal representations as mice adapt to high expected uncertainty of reward location.

### Acetylcholine release and neuronal plasticity in response to unexpected uncertainty

To examine how cholinergic signalling relates to dRSC plasticity under conditions of unexpected uncertainty, we measured GRAB-ACh4l or jGCaMP8m signals during relocation of the reward zone within a familiar visual environment, from a narrow (10 cm wide) reward zone 1 m along the track to a distinct narrow reward zone 2 m along the track (Fig. 4A). Relocation of the reward zone decreased anticipatory licking and deceleration behaviors, which partially recovered as mice gradually adapted their behavior over several laps to anticipate the new reward location (Fig. 1F). No increase in acetylcholine release was observed after the reward zone switch and although there was a small, inconsistent elevation in acetylcholine release immediately after the switch when mice had low velocity, this was not above random lap-by-lap fluctuations observed during the baseline. Unlike novel environment (Fig. 2) or expected uncertainty (Fig. 3) conditions, this manipulation did not significantly modulate acetylcholine release at early or late phases after the switch (Fig. 4B-E; Table S1). Nevertheless, relocating the reward zone elicited a rapid increase in pupil area that remained elevated throughout the remainder of the recording session (Fig. 4C-E; Table S2), indicating that mice quickly detected the change in reward location as a salient perturbation consistent with an experience of unexpected uncertainty.

Relocation of the reward zone did not induce widespread remapping of dRSC positional representations (Fig. 4F). A substantial proportion of cells (39.5±3.8%) remained positionally stable (Fig. 4G), significantly exceeding the level predicted under global remapping (Fig. 4H). In contrast, RCCs (19.0 ± 4.6% of cells) robustly tracked the new reward location (Fig. 4I-J) shifting their firing fields towards the end of the track in tandem with the relocated reward zone (Fig. 4F-J). Of these RCCs, 8.0 ± 2.3% of total cells were active in close proximity (≤25 cm) to the reward zone, while 11 ± 3.1% were not close to the reward zone (>25 cm) but exhibited stable activity in the reward reference frame (Fig. 4I,J). The shift of RCCs firing location by a similar distance to the reward location is similar to that reported for hippocampal CA1 ^37,42–44^. Together, these findings indicate that adaptations in dRSC neuronal representations in response to unexpected changes in reward location occur within a stable spatial reference frame in the absence of increased acetylcholine release.

**Figure 4.**
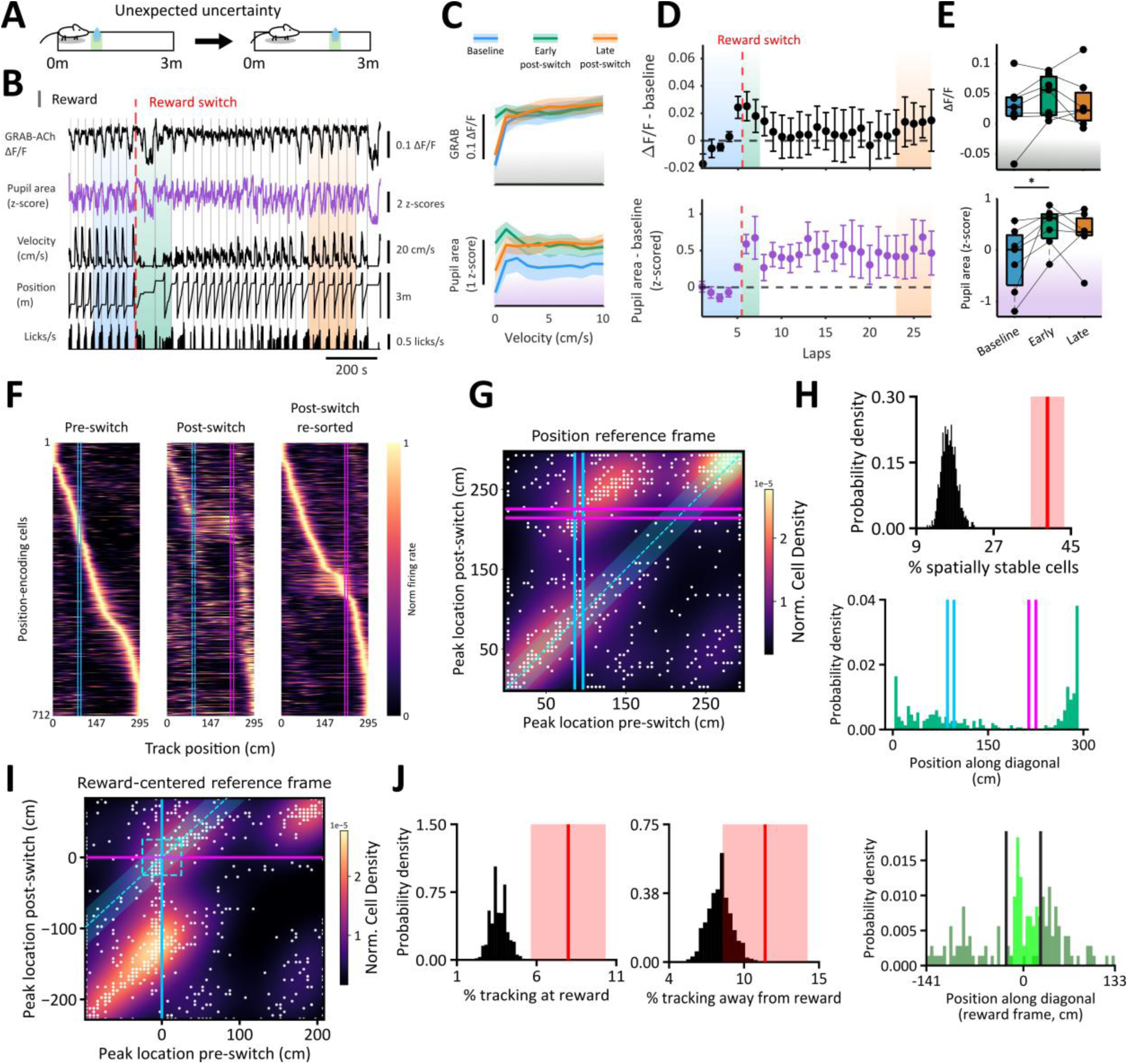
Acetylcholine release and neuronal representations in response to unexpected uncertainty. (A) Reward switch in which a narrow proximal (10 cm) reward zone transitioned to a narrow distal reward zone. (B) Representative recording showing GRAB-ACh4l fluorescence (ΔF/F), pupil area (z-scored), running velocity (cm/s), track position (m), and licking rate (licks/s). Red dashed line indicates the reward switch. (C) ΔF/F (top) and pupil area (bottom) binned by running velocity (1 cm/s bins). Separate lines indicate baseline (blue), early post-switch (green), and late post-switch (orange) epochs. N = 7. (D) Lap-by-lap baseline-subtracted ΔF/F (top) and pupil area (bottom) aligned to the reward switch (red dashed line). Shaded regions denote baseline (blue), early post-switch (green), and late post-switch (orange) epochs. Dashed lines indicate zero. (E) Mean ΔF/F (top) and pupil area (bottom) across behavioural epochs (baseline, early, and late). Repeated-measures ANOVA with Greenhouse–Geisser correction, followed by Dunnett-adjusted post hoc comparisons against baseline (\**p* < 0.05). (F) Normalized spatial activity maps for PCCs. Left, pre-switch activity maps sorted by peak firing location. Middle, post-switch maps for the same neurons aligned to pre-switch ordering. Right, post-switch maps reordered by peak location. Pre- and post-switch reward locations are indicated with cyan and magenta lines, respectively. (G) Galaxy plot showing the relationship between pre- and post-switch peak firing locations. White dots represent individual neurons; colour indicates population density (KDE). Cyan dashed diagonal and shaded band denote ±25 cm stability threshold. (H) Spatial shuffle analysis of stability across the reward switch. Top, null distribution with observed proportion of spatially stable cells indicated in red. Bottom, probability density of neurons along the diagonal axis, with pre- and post-switch reward zones indicated. Number of spatially stable cells was significantly greater than chance (*p* = 0.001; one-sided permutation test). (I) Reward-referenced galaxy plot. Peak firing locations are expressed relative to reward position. Cyan dashed diagonal indicates reward-aligned tracking; reward location for each trial is centred at zero. (J) Reward-tracking analysis in the radial reward-referenced frame. Left, null distribution for cells tracking at the reward location; middle, null distribution for cells tracking away from reward; right, distribution of neurons along the diagonal axis in a radial reward reference frame (±25 cm around reward indicated by black lines). Tracking at and away from the reward location were both significantly above chance (*p* = 0.0002 and *p* = 0.0062, respectively; one-sided permutation tests).

In addition to encountering unexpected uncertainty in otherwise stable environments, animals may also experience unexpected uncertainty against a background of already high expected uncertainty. To investigate how unexpected uncertainty interacts with expected uncertainty, we next performed a similar set of experiments where mice were initially trained under conditions of high expected uncertainty by varying the reward location within a broad (100 cm) reward zone centred 1 m along the track (Fig. 5A). Acetylcholine release is predicted to be high in this initial condition of elevated expected uncertainty (Fig. 3). We found that, similar to the condition of initial low expected uncertainty, acetylcholine release showed an initial small, transient, and inconsistent increase in response to a permanent shift in reward location to a distinct narrow (10 cm) zone 2 m along the track. However, in contrast to this manipulation from an initial state of low expected uncertainty (Fig. 4), acetylcholine release slowly increased over the course of the session (Fig. 5B-E; Table S1), although this was variable across mice. In parallel, the level of arousal as measured by pupil area also increased at a comparable rate (Fig 5B-E; Table S2).

Consistent with unexpected uncertainty from a background of low expected uncertainty, changing the reward zone from a broad zone 1 m along the track to a narrow zone 2 m along the track again did not produce widespread remapping of dRSC positional representations, with 47.4 ± 4.1% of cells remaining positionally stable (Fig. 5F-H). RCCs representing 16.3 ± 6.3% of total cells also robustly tracked the new reward location (Fig. 5I-J), shifting their firing fields towards the end of the track in line with the relocated reward zone (Fig. 5F-J). Of these RCCs, 7.1 ± 3.2% of the total cells were in close proximity (≤25 cm) to the reward zone.

Taken together, these findings show that dRSC representations adapt in a grossly similar manner when the reward zone is moved abruptly to induce unexpected uncertainty, regardless of whether the initial state is one of low or high expected uncertainty. However, the levels of acetylcholine release are higher when that manipulation occurs from a starting condition of high expected uncertainty.

**Figure 5.**
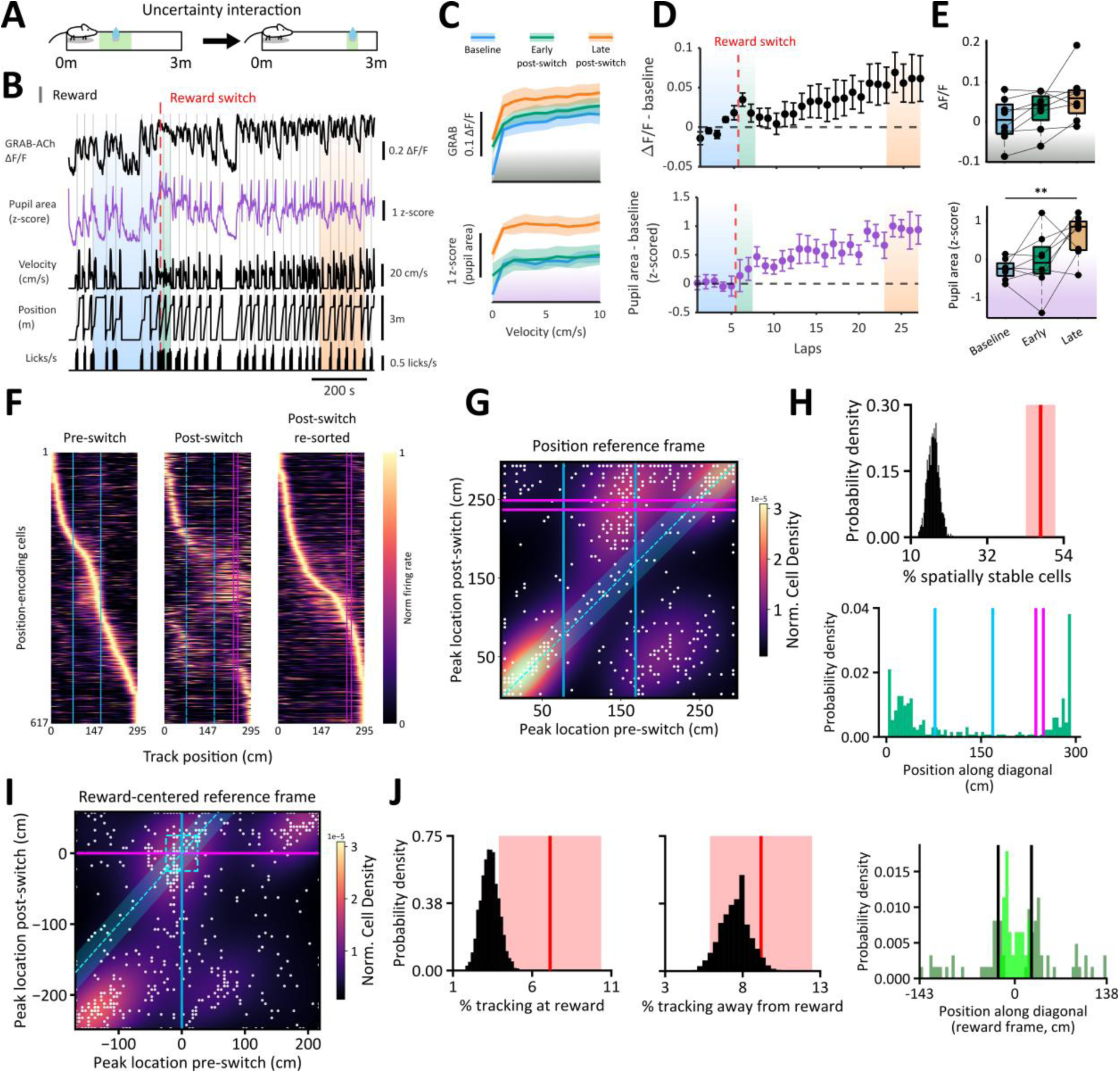
Acetylcholine release and neuronal representations in response to unexpected uncertainty from a baseline of high expected uncertainty. (A) Reward switch in which a broad proximal (1 m) reward zone transitioned to a narrow distal (10 cm) reward zone. (B) Representative recording showing GRAB-ACh4l fluorescence (ΔF/F), pupil area (z-scored), running velocity (cm/s), track position (m), and licking rate (licks/s). Red dashed line indicates the reward switch. (C) ΔF/F (top) and pupil area (bottom) binned by running velocity (1 cm/s bins). Separate lines indicate baseline (blue), early post-switch (green), and late post-switch (orange) epochs. N = 8. (D) Lap-by-lap baseline-subtracted ΔF/F (top) and pupil area (bottom) aligned to the reward switch (red dashed line). Shaded regions denote baseline (blue), early post-switch (green), and late post-switch (orange) epochs. Dashed lines indicate zero. (E) Mean ΔF/F (top) and pupil area (bottom) across behavioural epochs (baseline, early, and late). Repeated-measures ANOVA with Greenhouse–Geisser correction was used to assess main effects of epoch, followed by Dunnett-adjusted post hoc comparisons testing early and late epochs against baseline (\*\**p* < 0.01). (F) Normalized spatial activity maps for PCCs. Left, pre-switch activity maps sorted by peak firing location. Middle, post-switch maps for the same neurons aligned to pre-switch ordering. Right, post-switch maps reordered by peak location. Pre- and post-switch reward locations are indicated (cyan and magenta lines, respectively). (G) Galaxy plot showing the relationship between pre- and post-switch peak firing locations. White dots represent individual neurons; colour indicates population density (KDE). Cyan dashed diagonal and shaded band denote ±25 cm stability threshold. (H) Spatial shuffle analysis of stability across the reward switch. Top, null distribution with observed proportion of spatially stable cells indicated in red. Bottom, probability density of neurons along the diagonal axis, with pre- and post-switch reward zones indicated. Stability was significantly greater than chance (*p* < 0.001; one-sided permutation test). (I) Reward-referenced galaxy plot. Peak firing locations are expressed relative to reward position. Cyan dashed diagonal indicates reward-aligned tracking; reward location is centred at zero. (J) Reward-tracking analysis in the radial reward-referenced frame. Left, null distribution for cells tracking at the reward location; middle, null distribution for cells tracking away from reward; right, distribution of neurons along the diagonal axis (±25 cm around reward shown as black lines). Tracking at and away from the reward location were both significantly above chance (*p* = 0.0002 and *p* = 0.0304, respectively; one-sided permutation tests).

### Comparison of learning rates between conditions of uncertainty

To investigate the relationship between acetylcholine and neuronal plasticity we next compared the levels of acetylcholine release between conditions of uncertainty, with corresponding neuronal plasticity measured by the proportions of cells that adapt to environmental changes and the speed of neuronal and behavioral adaptations.

Acetylcholine release increased during transition from low to high expected uncertainty and did not increase when unexpected uncertainty was induced by moving the reward location between discrete low uncertainty locations (Figs. 3E,4E). Intermediate levels of acetylcholine release were found when unexpected uncertainty was induced from a condition of high expected uncertainty (when acetylcholine release was presumably initially high) and subsequently increased after the reward location switch in some mice but not others (Fig. 5E). As a reference point, exposure to a novel environment that is predicted to incur high levels of both unexpected and expected uncertainty induced high levels of acetylcholine release (Fig. 2E).

The differences in acetylcholine release suggest differences in neuronal plasticity under the various conditions of uncertainty. Neuronal plasticity can be measured by the speed of representational adaptations, which we analysed using a drift diffusion model of the neuronal population that tracked the representation as it adapted to the new environmental conditions after a switch ^45–47^. Lap-by-lap cross correlations of the population neuronal activity showed that representations changed abruptly after unexpected uncertainty or exposure to a novel environment but changed less when expected uncertainty increased (Fig. 6A,B). However, analysis of the neuronal activity manifold revealed that the pathway taken to a new stabilised representation was less direct when expected uncertainty increased in comparison to induction of unexpected uncertainty or presentation of a novel environment (Fig. 6A,D; Table S3). Subsequently, analysis of the lap-by-lap recurrency of the manifold revealed the number of laps required to reach a new stable representation was correspondingly low for increased expected uncertainty despite the relatively large distance of travel for the representation (Fig. 6E, S5; Table S4). The most direct comparison for this time to reach stability was between conditions of unexpected uncertainty where the initial condition was either low or high expected uncertainty. Here we found a significantly faster time to reach stability when unexpected uncertainty was preceded by a period of high expected uncertainty (Uncertainty interaction, Fig. 6E, S5; Table S4).

This faster stabilisation of neuronal representation was mirrored by a faster behavioral adaptation measured by the velocity and licking activity and analysed in the same drift diffusion model (Fig. 6C,E). Although unexpected uncertainty and novel environment switches produced stereotypical behavior as the mice learnt the new reward location, this took many laps to stabilise. In contrast, the expected uncertainty and uncertainty interaction switches had less stereotypical behavior after the switches, but the behavior stabilised in fewer laps. This was captured by the recurrency analysis (Fig. 6E, S5; Table S4). Importantly, acetylcholine levels during late post-switch periods were negatively correlated with the number of laps required for behavior to stabilise (Fig. 6F). Thus, increased acetylcholine release found in high expected uncertainty occurred alongside faster adaptations in behavior and neuronal representations under conditions of unexpected uncertainty.

**Figure 6.**
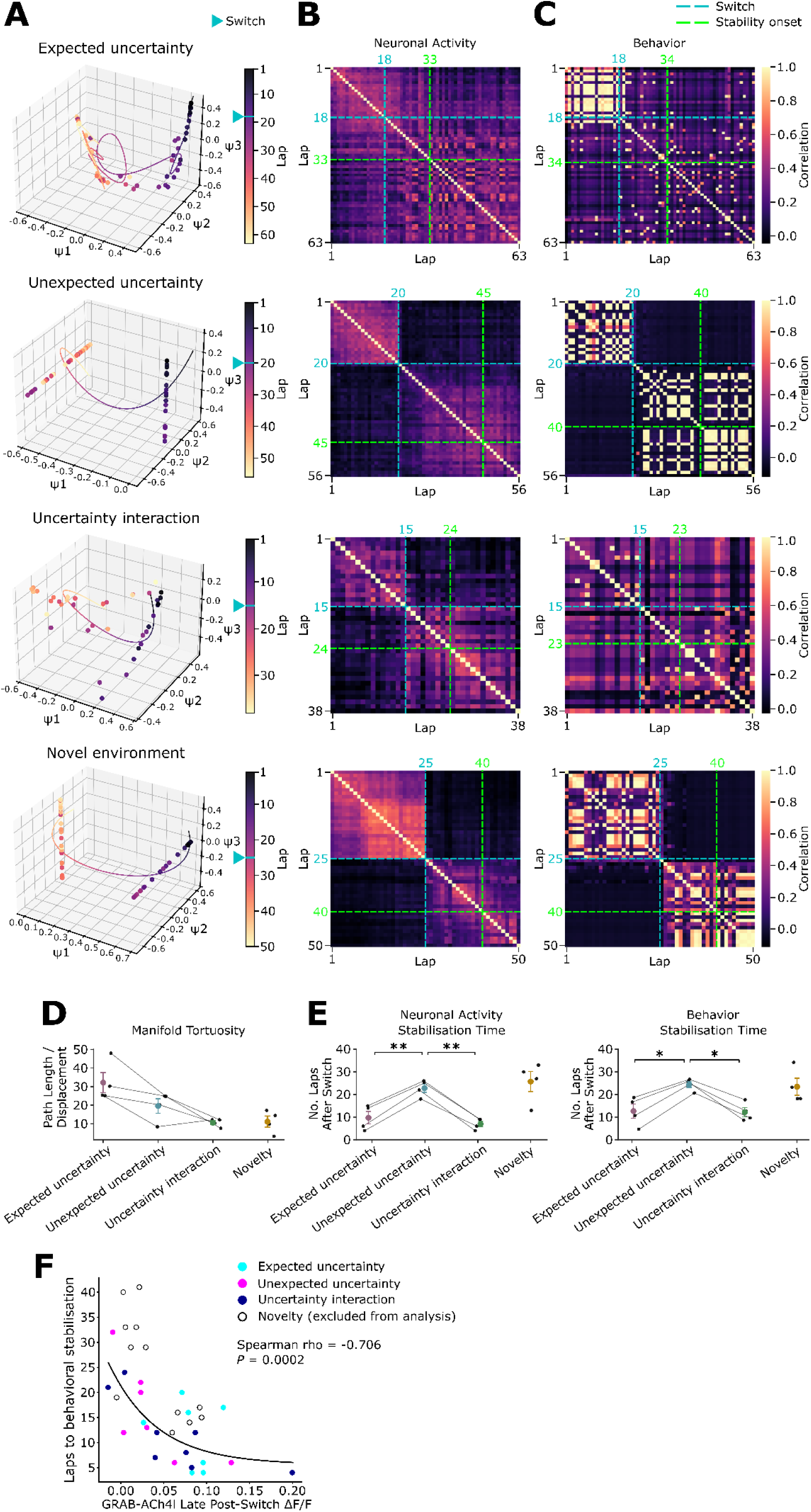
Faster adaptations to behavior and neuronal representation in uncertainty conditions with high acetylcholine release. (A) Example diffusion maps illustrating the evolution of neuronal representations across switch conditions. Each dot corresponds to the eigenprojection of neuronal population activity from a single lap of the virtual track, coloured by lap number. The accompanying curve shows diffusion distance between consecutive laps, where smaller distances indicate greater representational similarity. Blue arrows mark the lap on which the environmental manipulation (Switch) occurred. (B) Population correlation vector heatmaps for neuronal representations across switch conditions, from which the manifolds in the corresponding rows of panel A are derived. Dashed blue lines indicate the lap of the switch; dashed green lines mark the first post-switch lap at which the neuronal representation was classified as stable. (C) Corresponding behavioral population correlation vector heatmaps across switch conditions, based on combined running velocity and licking behavior. Dashed lines follow the conventions in panel B, with the green line indicating the first post-switch lap at which behaviour was classified as stable. (D) Manifold tortuosity across switch conditions (repeated-measures ANOVA; *p* > 0.05). (E) Onset of stability for each switch condition in neuronal representation (left), and behaviour (right) (repeated-measures ANOVA; Bonferroni-corrected paired t-tests; \**p* < 0.05; \*\**p* < 0.01). (F) Mean GRAB-ACh4l ΔF/F of the late post-switch epoch plotted against the onset of behavioral stability (Spearman ρ = -0.706, *p* = 0.0002; single exponential curve fit). To enable meaningful statistical comparison between conditions where uncertainty was manipulated by altering reward predictability within an otherwise stable environment, rather than through wholesale environmental change, the Novel Environment condition was excluded from statistical analyses in panels D-F.

## Discussion

The relationship between acetylcholine release, plasticity, and uncertainty has remained largely untested empirically since it was first proposed that acetylcholine is associated with increased expected uncertainty and enhanced plasticity ^1,6,7^. This picture has been further complicated by findings linking acetylcholine to surprise ^12,48^, raising the possibility that acetylcholine may also, or instead, signal unexpected uncertainty. Here we provide data that disentangles these roles, demonstrating that tonic acetylcholine signals expected uncertainty rather than unexpected uncertainty per se. We suggest a revised model where acetylcholine tracks changes in the reliability of external cues, marking transitions in the environmental state that require integration into neuronal representations of context.

### Acetylcholine signals a state change in expected uncertainty

We examined acetylcholine release in the dRSC in response to manipulations of environmental context designed to selectively alter expected or unexpected uncertainty, either independently or in combination. Acetylcholine release was associated with increased expected uncertainty induced by reducing the reliability of reward location around a fixed position within a familiar environment (Fig. 3). In contrast, acetylcholine release was not observed when unexpected uncertainty was introduced in an otherwise stable context by abruptly relocating the reward to a different region of a familiar environment (Fig. 4). Notably, acetylcholine release was observed following exposure to a novel environment (Fig. 2), a manipulation that constitutes a large contextual shift predicted to increase both unexpected uncertainty and, importantly, expected uncertainty as new contingencies are learned. Thus, these initial findings broadly support theoretical models proposing that acetylcholine signals expected, rather than unexpected, uncertainty ^1,7^.

We also observed increased acetylcholine release when reward delivery shifted from an unreliable, broad zone to a reliable, narrow zone in a distinct region of the track – a manipulation that introduced unexpected uncertainty against a background of high expected uncertainty (Fig. 5). However, because acetylcholine release was selectively modulated by manipulations of expected uncertainty but not unexpected uncertainty in other experiments, we consider it unlikely that unexpected uncertainty accounted for the observed increase in cholinergic tone. Instead, we propose that this increase in acetylcholine release reflects a transition between distinct states of expected uncertainty, specifically, a shift from high expected uncertainty (broad reward zone) to low expected uncertainty (narrow reward zone).

The interpretation that acetylcholine is sensitive to state transitions in reward reliability is consistent with emerging work showing that acetylcholine release scales with uncertainty of reward outcome and can predict adjustment of behavioural strategy ^49^. However, our data further indicate that, at least in dRSC, acetylcholine release does not simply signal the absolute magnitude of expected uncertainty ^1^. Rather, acetylcholine appears to signal changes in reward predictability that reflect transitions in the expected uncertainty state – whether from low to high (Fig. 3) or from high to low (Fig. 5).

The sensitivity of acetylcholine release to bidirectional transitions in expected uncertainty resonates with evidence that basal forebrain cholinergic neurons respond to reinforcement signals of both positive and negative valence ^12^. These cholinergic populations project widely throughout the forebrain, including to dRSC ^17,50^, raising the possibility that changes in reward predictability drive activity within these circuits that underpin acetylcholine release observed in the present study.

Surprising, or unexpected, reinforcement can elicit phasic activation of basal forebrain cholinergic neurons, predominantly among those exhibiting a burst-firing phenotype ^12,48^. Although we did not observe significant modulation of acetylcholine release by unexpected uncertainty, we did not explicitly characterise phasic release. This was because acetylcholine release is tightly associated with movement velocity (Fig. S2) ^38,51,52^, which made it difficult to identify phasic events that could be robustly dissociated from transitions in velocity. Instead, we focussed on tonic signals sustained over multiple laps of track exploration. Tonic and phasic acetylcholine are thought to subserve distinct functions, with tonic release linked to arousal and attentional engagement, and phasic release associated with cue detection ^52,53^. Thus, our findings do not preclude phasic cholinergic responses to surprising reinforcement events, such as unanticipated rewards delivered under conditions of high expected, or unexpected, uncertainty. Rather, they support a model in which transitions in expected uncertainty primarily modulate tonic acetylcholine levels.

Because tonic acetylcholine is positively modulated by attention ^8,52^, and acetylcholine dynamics closely tracked changes in pupil area as a proxy for attentional state ^54^, acetylcholine release driven by changes in expected uncertainty may reflect the orientation of animals’ attention toward shifting reward contingencies. Importantly, however, acetylcholine did not function as a general attentional or reward prediction signal ^55^, as unexpected uncertainty elicited pupil dilation without corresponding modulation of acetylcholine release (Fig, 3).

### Acetylcholine is associated with rapid adaptation of dRSC representations and animal behaviour during expected uncertainty transitions

Neuronal representations in dRSC are sensitive to contextual changes ^14,32^. We therefore examined the plasticity of dRSC representations to manipulations of expected and unexpected uncertainty on the virtual linear track. Consistent with prior studies ^19,27,28,32^, dRSC populations robustly encoded positional and reward-related information. Putative position-encoding neurons also over-represented the start and end regions of the track, reminiscent of trial onset-encoding by dRSC neurons described previously ^27,30^. However, because trial start and end locations remained constant across laps, we did not attempt to disambiguate whether these populations primarily represented track position or aspects of trial structure.

We identified two populations of reward-encoding neurons: cells active near the reward location (‘at reward’ RCCs) and cells that tracked shifts in reward location from positions distal to the reward site (‘away from reward’ RCCs) ^44^. In our URTask, reward location effectively functions as a hidden landmark that is stable or variable, depending on task contingency. Notably, the behaviour of ‘away from reward’ RCCs was analogous to previously characterised dRSC populations anchored to reference frames defined by visual or tactile landmarks ^27,56^, suggesting that dRSC representations integrate salient environmental cues across both sensory and non-sensory domains.

Representational plasticity was induced by changing the degree of expected uncertainty, with fewer PCCs and a concomitant increase in recruited RCCs when expected uncertainty was initially increased (Fig. 3). The co-occurrence of elevated acetylcholine release and reorganisation of dRSC representations supports the hypothesis that acetylcholine facilitates plasticity under conditions of increased expected uncertainty ^1^.

We further assessed neuronal plasticity by quantifying the rate at which dRSC representations adapted following manipulations of expected or unexpected uncertainty (Fig. 6). Both representational updating and behavioral adaptation to the new reward contingency occurred more rapidly during transitions in expected uncertainty – conditions that were associated with elevated acetylcholine release. This might suggest that both dRSC representations and behaviour adapt more slowly in response to unexpected uncertainty. However, we consider this unlikely. In the uncertainty interaction condition, unexpected uncertainty was introduced against a background of high expected uncertainty, and both neural representations and behaviour adapted more rapidly than when unexpected uncertainty was introduced in isolation. This indicates that adaptation rates cannot be attributed solely to unexpected uncertainty. Thus, we propose that acetylcholine release in dRSC is important for both neuronal plasticity and adaptive behavior during uncertainty.

### Acetylcholine may support sensory-driven updating of contextual representations in dRSC

A proposed role of acetylcholine is to suppress reliance on expectation-driven knowledge while prioritising sensory information about the current state of the environment ^6,57^. Our findings are consistent with this view. When the level of expected uncertainty changes, previously learned contingencies may no longer be sufficient to guide behavior, requiring internal representations to be updated to support adaptative responses ^1^. Under these conditions, animals must shift from knowledge-guided behavior to actively sampling the reliability of environmental cues and incorporating this sensory evidence into contextual representations of the environment’s structure.

Consistent with this model, we found that increased expected uncertainty was accompanied by a gradual elevation of acetylcholine release across successive laps of track exploration. This pattern is consistent with tonic acetylcholine facilitating the prioritisation of sensory information related to the altered reward contingency. In parallel, increasing the variability of reward location induced remapping of dRSC representations, characterised by the recruitment of a greater proportion of reward-encoding neurons. Together, these findings suggest that acetylcholine supports learning under conditions of high expected uncertainty by promoting the integration of sensory-derived information into updated representations of the environment.

### Potential mechanisms of dRSC plasticity during uncertainty

The shift in the proportions of dRSC neurons encoding position and reward information parallels recent findings in hippocampal CA1. In CA1, moving the reward location changes the balance of place cells tuned to reward versus position reference frames ^42,44^, and this prioritisation of reward-encoding aligns with changes in the weights of synaptic inputs, regulated by the salience – or uncertainty – associated with the reward location. This process is proposed to depend on synaptic plasticity mechanisms, and specifically, behavioral timescale synaptic plasticity (BTSP) ^42^.

In CA1, position and reward information are thought to be transmitted from the medial and lateral entorhinal cortex, respectively ^58^. Notably, both medial and lateral entorhinal cortices project to dRSC ^59–61^, suggesting a potentially analogous route through which spatial and reward-related signals could converge to drive updating of dRSC representations. Alternatively, such information could be conveyed through CA1, which projects indirectly to dRSC via the subiculum, presubiculum, and parasubiculum ^61,62^ Supporting this possibility, lesion studies show that hippocampal input is important for remapping dRSC positional representations in novel environments ^35^, suggesting that the hippocampus may also contribute to dRSC remapping under conditions of uncertainty. Crucially, if dRSC representations can be similarly modified by BTSP, as emerging evidence suggests ^63^, then similar plasticity mechanisms could underpin the shifts in relative proportions of position- and reward-encoding neurons observed in our experiments.

In addition to prioritising sensory input ^57^, acetylcholine can enhance or “gate” the induction and expression of synaptic plasticity. In both cortical and hippocampal pyramidal neurons, acetylcholine enhances dendritic and spine excitability, amplifying calcium signalling that drives synaptic plasticity ^64–68^. Importantly, this creates the conditions required for large plateau potentials that induce BTSP ^69–71^. Acetylcholine also reconfigures local inhibitory circuits via its actions on interneurons expressing specific nicotinic and muscarinic receptor subtypes ^57,72–75^, producing a net disinhibitory effect on pyramidal neuron dendrites that further facilitates synaptic plasticity ^76,77^. Thus, we speculate that acetylcholine may support multiple complementary forms of plasticity within dRSC, promoting representational updating during transitions in expected uncertainty by enhancing sensory-driven input and enabling synaptic modifications needed to reconfigure neuronal representations.

## Concluding remarks

Our findings identify the dRSC as a key node in the neural processing of uncertainty and link cholinergic signalling to representational plasticity within this region. We show that acetylcholine release in dRSC increases when changes in reward predictability signal transitions in expected uncertainty, and that this coincides with the updating of neuronal representations of the environment. Together, these results support a model in which acetylcholine enables sensory evidence about changing environmental contingencies to be incorporated into internal representations. More broadly, this process may allow the flexible updating of cortical networks to support adaptive behaviour in dynamic environments.

## Methods

### Animals

All procedures were carried out in accordance with the UK Animals (Scientific Procedures) Act 1986 and were approved by the University of Bristol Animal Welfare and Ethical Review Board. 31 adult C57BL/6J mice (17 males, 14 females), aged 8-12 weeks at the time of surgery, were used across experiments. Mice were obtained from Charles River. Prior to behavioural training, mice were group housed in standard cages under a 12-hour light/dark cycle (lights off at 09:00, on at 21:00) with *ad libitum* access to food and water. During behavioural training, mice were singly housed and placed on a water scheduling regimen to promote task engagement, receiving 0.0375 ml water per gram of body weight per day, and maintained at or above 85% of their *ad libitum* feeding weight, calculated from the average of 3-5 days prior to water scheduling. Water was delivered either manually or via a WaterR device (https://github.com/DodsonLab/WaterR). Running wheels were provided in the home cage for environmental enrichment. Housing rooms were maintained at an ambient temperature of 21 °C and 45% relative humidity. The health and weight of animals were monitored daily throughout water scheduling.

### Surgical procedures

Mice were anaesthetised with isoflurane in oxygen (4% induction, 1-2% maintenance), their scalp was shaved, and they were placed in a rodent stereotaxic frame. Carprofen (5 mg/kg) was administered for analgesia, and the anti-inflammatory dexamethasone (4 mg/kg) was given to mice receiving a cranial window implant. Both drugs were administered via subcutaneous injection at the start of surgery. Lacri-Lube ointment (Allergan Inc.) was applied to the eyes to prevent drying. Body temperature was maintained at ∼37°C throughout the procedure using a feedback-controlled heat pad. Sterile saline (10 µL/g body weight, subcutaneous) was administered every 90 minutes to prevent dehydration.

The scalp was disinfected with chlorhexidine and incised, and the dorsal skull surface was cleared of connective tissue. The loose edges of the skin were fixed to the skull using cyanoacrylate glue (Vetbond). The exposed skull was levelled (within 0.1 mm in the dorsoventral axis) and scored using a sterile scalpel blade to improve the adhesion of subsequent cement layers. A 3 mm circular craniotomy was drilled, centred over the midline at 2.2 mm caudal to Bregma. During drilling, the skull was irrigated with chilled artificial cerebrospinal fluid (aCSF; in mM: 150 NaCl, 2.5 KCl, 10 HEPES, 1 CaCl2, 1 MgCl2; pH 7.3) to minimise heating.

Adeno-associated viral vectors (AAV) encoding GRAB-ACh4l, GRAB-ACh-mut ^39^, or jGCaMP8m ^40^ (**Table 1**) were injected at four sites (200 nL per site, two injections per hemisphere) using a 35G NanoFil needle (World Precision Instruments). An initial injection was made at ∼2.0 mm caudal to Bregma and 300–600 µm lateral to the midline, with subsequent injection sites arranged to form a square approximately 0.65 mm in anterior-posterior and mediolateral extent. Coordinates were adjusted slightly between animals to avoid surface vasculature. The injection depth was 0.3 mm from the brain surface. AAV was infused at 50 nL/min, with the needle left in place for 5–10 min post-injection to allow for diffusion. The needle was then slowly withdrawn at 10 µm/s to minimise backflow.

**Table 1.** AAV vectors used in the study.

| Name | Working titre (vg/ml) | Supplier | Product Code |
| --- | --- | --- | --- |
| AAV9-hSyn-GRAB-ACh4l (ACh4L) | $3.3 \times 10^{12}$ | BrainVTA | PT-8231 |
| AAV9-hSyn-GRAB-ACh3.0mut | $3.5 \times 10^{12}$ | BrainVTA | PT-1336 |
| AAV9-hSyn-jGCaMP8m-WPRE | $2.5 \times 10^{12}$ | Addgene | 162375-AAV9 |

A custom cranial window was affixed over the craniotomy using cyanoacrylate glue (VetBond or Loctite). Each window was composed of one 5 mm and two 3 mm circular coverslips (#1 thickness; CS-3R and CS-5R; Warner Instruments) stacked and bonded with Norland Optical Adhesive #81 (Edmund Optics). This configuration allowed the larger 5 mm coverslip to adhere to the skull while the smaller coverslips occupied the excised skull volume to limit bone regrowth and improve subsequent imaging stability ^27^.

A custom head bar with an integrated imaging aperture and light block interface ring was affixed to the skull using C&B Super-Bond (Sun Medical), followed by a layer of gentamicin-impregnated bone cement (DePuy Synthes) mixed with activated charcoal to reduce light leakage during imaging. Mice used for non-imaging behavioural experiments received surgical implantation of a linear head bar without accompanying AAV injection or implantation of a cranial window.

### VR behavioural task

The VR setup was similar to previously described systems ^58^. Five flat-screen monitors were arranged vertically in a half-octagon, positioned ∼38 cm from the mouse, and covering ∼220° horizontally and ∼68° vertically of the mouse’s visual field. Visual stimuli were rendered at 5,827 × 1,920 resolution using the ViRMEn engine ^78^.

Mice were head-fixed and trained to run along a 3-metre-long virtual linear corridor. Each side of the corridor featured both proximal (e.g., vertical poles) and distal (e.g., towers or distant landmarks) visual cues, along with patterned walls, to provide robust optic flow and spatial cues to aid navigation. Mice traversed the VR environment by running on a custom polystyrene treadmill (∼20 cm diameter, ∼7.5 cm width). Locomotion was tracked by a rotary encoder (US Digital EM2-1-5000-I) attached to the treadmill axle, continuously measuring angular displacement which was translated to VR movement. 10% sucrose (4 µL) rewards were delivered when mice entered a designated reward zone, via a lick spout controlled by a solenoid pinch valve. Licking of the spout was detected by a capacitive touch sensor. When mice reached the end of the track, the visual display went dark for a 3 s inter-trial interval, after which the mouse was automatically returned to the start of the corridor to commence the next trial. Behavioural data streams – including VR position, movement velocity, licking, and reward delivery – were recorded using ViRMEn (> 100 Hz sampling rate). Behavioural monitoring was also performed online using a Micro1401 analogue-to-digital data acquisition interface and Spike2 software (CED, Cambridge, UK).

### Two-photon imaging

Two-photon imaging was performed using a custom-built microscope (Cosys Ltd.) equipped with a resonant galvanometric scanner (ThorLabs) and a Ti:Sapphire laser (Mai Tai, Spectra-Physics) tuned to 920 nm for excitation of GRAB-ACh4l, GRAB-ACh-mut, and jGCaMP8m. Laser power at the sample was 20-50 mW. Fluorescence emission was collected through a 16× water-immersion objective (Nikon, CFI LWD Plan Fluorite, 0.8 NA, 3 mm WD), bandpass filtered using a FF01-510/84 filter (Semrock), and detected with a GaAsP photomultiplier tube (H10770PA-40, Hamamatsu Photonics). Light contamination from the VR screens was minimised by a custom 3D-printed light-shield affixed to a 3D-printed interface ring mounted on the head bar. Black nylon material (from a balloon) was positioned between the light-shield and objective to fully occlude ambient light.

Image acquisition was controlled using ScanImage (v5.7 R1) software (MBF Bioscience) running in MATLAB (MathWorks), which was synchronised with the VR environment by transmitting a frame synchronisation pulse from the VR computer to the microscope data acquisition card (National Instruments). Time series videos (∼15,000-60,000 frames/session; 512 × 512 pixels/frame) were collected at ∼29.8 Hz. The field of view was approximately 850 × 850 µm, and most imaging sessions were performed at 2× digital zoom. GRAB-ACh imaging was performed in layer II/III of dRSC, ∼120–150 µm from the brain surface; layer I was readily identified by its sparse somatic density and prominent neuropil, allowing reliable placement of the imaging plane ventral to this level in layer II/III ^62^. For jGCaMP8m imaging, neurons in layer II/III were longitudinally tracked across days by manually aligning fields of view using reference images of blood vessels and cell morphology.

### Pupil imaging

Pupil area was recorded during VR behaviour using a monochrome industrial camera (DMK 37BUX273, The Imaging Source) positioned to capture the left eye of the mouse. The camera was angled laterally to avoid obstructing the mouse’s central visual field. The pupil was indirectly illuminated through tissue scattering of the two-photon imaging laser. Pupil image time series were synchronised with behavioural and two-photon imaging data via pulses sent from the VR software to a Micro1401 analogue-to-digital interface running Spike2 software. Spike2 generated a 25 Hz digital pulse that triggered each camera frame. Pupil videos were processed offline in MATLAB using the MEYE toolbox ^79^.

### Pupil analysis

Pupil area was extracted from the image time series using a deep learning-based segmentation model (MEYE) ^79^ with a resampling frequency of 23 Hz. Frame timestamps were obtained from image metadata, referenced to the experiment start time. For each frame, a binary pupil mask was generated and cleaned using morphological opening (using a circular structuring element of 5-pixel radius) to remove noise pixels. Pupil area was calculated from the pupil mask, and values were temporally upsampled via linear interpolation to match the two-photon imaging timebase, ensuring temporal alignment between pupil and neural signals. Finally, pupil area time series were smoothed using a Savitzky-Golay filter (polynomial order 3) with 2-second window size.

### GRAB-ACh4l image processing

For in vivo GRAB-ACh4l and GRAB-ACh-mut recordings, image time series were spatially downsampled from 512×512 to 256×256 pixels using nearest-neighbour interpolation. Motion correction was performed with a subpixel registration method based on Discrete Fourier Transformation ^80^, with each frame aligned to a reference image generated from the average projection of the first second (30 frames). GRAB-ACh fluorescence signals were isolated by defining a region of interest (ROI) spanning the top ∼80% of pixel intensities in the mean projection image, thereby excluding regions of poor GRAB-ACh expression or vasculature.

A background signal was estimated from a manually drawn vascular mask and subtracted from the main ROI signal. This procedure preserved slow fluctuations in the GRAB-ACh signal while correcting for noise. The background-subtracted signal was then normalized to the median fluorescence over the entire time series to produce a ΔF/F trace, which was subsequently smoothed using a Savitzky-Golay filter (polynomial order 3) with a 2 s window.

### Velocity-controlled analysis of GRAB-ACh4l

To control for the influence of running speed on GRAB-ACh4l signal, ΔF/F traces were binned according to running velocity (1 cm/s bins) up to 10 cm/s. Within each bin, ΔF/F values were averaged to generate velocity-controlled estimates of acetylcholine signalling.

### jGCaMP8m image processing

For calcium imaging, two-photon image series were processed using Suite2P ^81^. Motion correction was performed with rigid and non-rigid registration, using a 500-frame mean image as the reference template. Images were acquired at ∼29.8 Hz. Regions of interest (ROIs) corresponding to putative neurons were automatically detected using Suite2P’s segmentation algorithm, with the decay time constant (τ) set to 0.6.

Calcium fluorescence signals from each ROI were first corrected for neuropil contamination by subtracting the surrounding neuropil signal multiplied by a factor of 0.7, as provided by Suite2P. Slow fluctuations in the neuropil-corrected traces were removed using a sliding percentile baseline. For each time point, the 8th percentile of the fluorescence within a 60-second window was subtracted, and the resulting trace was normalized to yield ΔF/F₀. Significant fluorescence transients were then identified using a double-thresholding procedure: a transient was defined to start when the trace exceeded 2 standard deviations above baseline and to end when it returned below 0.5 standard deviations. Positive and negative transients were classified separately, and a histogram-based analysis of the ratio of negative-to-positive-going events across amplitudes and durations was used to determine thresholds that limited false positive events to <1%. Only transients meeting these criteria were preserved in the final trace, with all other values set to zero. These processed traces were used for all subsequent analyses.

### Cell registration across days

For analysis in Fig. S4, cell identities were tracked across sessions using the MATLAB implementation of CellReg ^82^. Spatial footprints obtained after motion correction and ROI extraction were aligned across sessions, and pairwise similarity matrices based on centroid distance and spatial correlation were used for probabilistic registration. Cells with match probabilities >0.5 were classified as the same cell across days, with only one-to-one matches retained. Registration quality was verified by visual inspection of matched footprints. The resulting registration matrix was used to follow cell properties across experimental conditions.

### Position and reward cell identification

To identify neurons with spatially selective activity, the spatial information (SI) of each cell’s calcium fluorescence across the virtual track was calculated ^83^. Fluorescence traces were first filtered to include only frames where the animal’s velocity exceeded 5 cm/s. For each cell, activity was binned by position along the track (60 bins spanning 0–300 cm), and a Gaussian smoothing (σ = 2 bins) was applied to both the binned activity and occupancy maps.

The spatial information of each cell was then computed as:

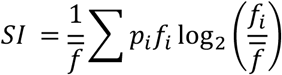

where *f_i_* is the mean fluorescence in bin *i, f̄* is the mean fluorescence across all bins, and *p_i_* is the probability of occupancy in bin (time spent in bin divided by total time). Higher SI values indicate that a cell’s activity is more strongly modulated by position.

To assess significance, the spatial information of each cell was compared against a shuffled distribution obtained by randomly circularly permuting the cell’s activity 1000 times relative to position. Cells with p < 0.05 (z-score calculated relative to the shuffled distribution) were classified as position-encoding cells. Activity maps for each cell were calculated by dividing the smoothed binned activity by the smoothed occupancy map, and normalized activity maps were obtained by dividing by the peak bin value.

Cross-validation (Fig. S4) was performed by calculating spatial information and significance separately for odd and even laps using the same procedures as the full dataset. Activity maps from odd laps were aligned according to the peak positions observed in the corresponding even-lap activity maps.

Spatially stable cells were defined as neurons whose field peak locations remained consistent across two conditions (i.e., pre vs. post switch, or odd vs. even laps). For each cell, the peak position of the normalized activity map was calculated in both conditions using identical spatial binning (59 bins across a 295 cm track). A cell was classified as spatially stable if the absolute difference between peak positions across the two conditions was ≤25 cm.

Reward-tracking cells were defined according to their spatial tuning relative to reward locations. A reward reference frame was generated by subtracting the reward location of each lap from all positions within that lap. Reward-aligned activity maps were then constructed, and cells were classified as reward-tracking *at* reward if their peak activity was located within ±25 cm of the reward in both pre- and post-switch sessions. Cells were classified as reward-tracking *away* from reward if their peak activity shifted with the reward location (within ±25 cm of the reward shift) but remained outside ±25 cm of the reward in both sessions.

### Rate maps

The calcium fluorescence traces were transformed into rate maps that represent the averaged cell activity in spatial bins across all laps for that entire session. The track was divided into 60 bins that were 5 cm long. The calcium traces were grouped into laps by identifying when the coordinate readout of the mouse reset to its minimum. The act of resetting the position of the mouse had the potential to result in laps that were impossibly brief (lasting for only a few sample frames). These laps were ignored. The cells were scaled using their respective peak activity, and the rate maps were normalized using the occupancy of each spatial bin. The rate maps were Gaussian smoothed across bins (σ = 2). The output of this pre-processing was a data structure of shape (*L, B, N*), where *L* is the number of laps, *B* is the number of bins, and *N* is the number of cells. All cells were analysed, regardless of spatial tuning.

### Lap-level population vector correlation

Rate maps were converted into a lap-by-lap population vector correlation (PVC) matrix, using the Pearson correlation. An entry in the matrix describes how similar a rate map at a given lap is to a rate map at another lap.

### Manifold analysis

To analyse the lap-by-lap activity, the PVC matrix was used to generate a diffusion map. A diffusion map is a nonlinear dimensionality reduction method that uses the local geometry of the input to embed it into Euclidean space ^84^. To build the diffusion map, the PVC matrix was used to derive a symmetrised nearest-neighbours distance graph *W* . This graph describes the local connectivity of lap pairs, instead of describing all pairwise distances. The number of neighbours was set to the square root of the number of laps multiplied by two. A Gaussian kernel (σ = 0.9 for representation, σ = 2 for behaviour, selected based on the median non-zero pairwise distance in the KNN graph, averaged across experiments) was applied to the graph to convert it into an affinity matrix *K*:

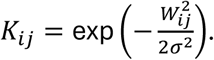

To mitigate the effect of non-uniform sampling density (for example, laps at the start and end of a session would have fewer neighbours), the affinity matrix was scaled using:

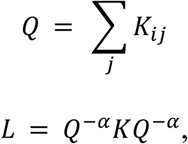

where α is a sampling density parameter. α was set to 1, which approximates the Laplace-Beltrami operator, resulting in a matrix that is unaffected by sampling density. It also reduces the sensitivity to changing the kernel bandwidth size. A Markov diffusion operator *P* was derived from this scaled adjacency matrix:

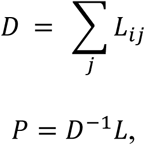

where the result *P* is a transition matrix of a random walk on the graph. In essence, *P* describes how likely a point on the manifold is to diffuse into another neighbourhood. *P* is eigendecomposed into its three leading eigenpairs. The spectral gaps of the eigenvalues of *P* were largest after the second or third eigenvalue. Three dimensions were chosen to capture as much of the dynamics as possible without introducing unnecessary noise. The eigenvectors must be converted into diffusion coordinates:

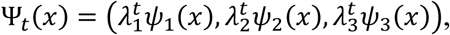

where Ψ*_t_*(*x*) are the 3D diffusion coordinates at a given discrete timestep *t,* λ is an eigenvalue, and *ψ* is an eigenvector. The diffusion distance between coordinates measures the similarity between them based on their connectivity. The Euclidean distance between points in the embedding space approximates the diffusion distance, and it is easier to compute and interpret geometrically. For this reason, the Euclidean distance was used to compare points on the manifold rather than the diffusion distance. The Euclidean distance a point in the embedding space moves from one lap to another describes the magnitude of change in population vector activity. To analyse the learning dynamics after a switch has occurred, the manifold was segmented into a pre-switch part and a post-switch part. The total distance covered by the post-switch manifold and the final displacement from its post-switch starting point were computed and used to calculate the tortuosity. The tortuosity is the ratio of the total distance covered to the final displacement and is a measure of both how efficient the path the manifold took was, as well as the amount of flickering in the map. A tortuosity equal to 1 means that the path taken was completely efficient, and that the manifold did not wander. A tortuosity equal to *T* can be interpreted as the manifold moving *T* times more than it needed to.

### Representational stability

Recurrence quantification analysis (RQA) was used to identify the point at which learning stabilised. RQA is a way of analysing nonlinear systems by quantifying how often a system tends to revisit previously visited states ^85,86^. Only the post-switch part of the manifold was considered, and it was used to compute the pairwise distance between all points in that embedding. Additionally, the manifold was upsampled using cubic spline interpolation to denoise the coarse signal. The upsample factor was tested with different values (1x, 2x, 4x, 8x), and the trend of RQA metrics was consistent across tests. A Heaviside step-function with a tolerance that was dynamically calibrated to the pairwise distance matrix was used to transform the distance matrix into a binary recurrence matrix *R*, where a value of 1 at *R_ij_* indicates that the manifold roughly visited the same area at lap *j* as it did at lap *i*. The tolerance was calibrated using the pairwise distance matrix and a target recurrence density of 10%. The recurrence rate, RR, measures the density of recurrence points where:

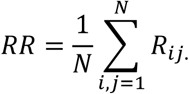

The *RR* is the percentage of times the system returns to a previously visited state. Points in *R* can form diagonal lines of ones. Many long diagonal lines of ones indicate a stable and deterministic system. By counting the number of persistent diagonal lines of ones, you can compute a metric *DET* ^87^:

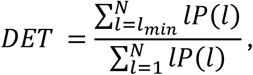

where *P*(*l*) is the frequency distribution of the lengths of these diagonal lines, and *l_min_* is the shortest diagonal line length considered (set to 2). Likewise, the percentage of recurrence points that form vertical lines is called laminarity and can be useful for identifying periodic oscillatory dynamics ^88^. Laminarity is taken to be:

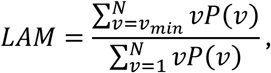

where *P*(*v*) is the frequency distribution of the lengths of these vertical lines, and *v_min_* is the shortest vertical line length considered (set to 2). These metrics were calculated using a sliding window on *R*, where the width of the window was equal to the number of laps divided by six, with a minimum width of three laps. After the windowed metrics were collected, they were combined into a composite stability score *S* that is the average of the three metrics and measures the stability of the system over the post-switch part of the session. A series of statistical tests were used to identify the point at which stability occurred. A Pettitt test is a non-parametric test for identifying the single most significant breakpoint in time series data ^89^. Using *S* as the input, a first candidate stability onset τ was identified with the Pettitt test, where τ is a lap number. If this breakpoint was significant (*p* ≤ 0.01), then this initial guess τ was investigated further. Otherwise, the maximum number of laps was returned instead. *S* was then segmented, looking from τ onwards. Another statistical test was used to assess whether the system was truly stable after τ. The Hamed-Rao test inspects whether time series data exhibits a significant trend in either direction. The Hamed-Rao test is a version of the Mann-Kendall test that has been adapted for autocorrelated data, and time series recordings of neurons are highly autocorrelated. If the Hamed-Rao test returned that the system after lap τ displayed no significant trend, and Sen’s slope was approximately zero, then τ was accepted as the onset of stability ^90,91^. If there was a significant trend, then τ was rejected, and another Pettitt test was conducted on *S* looking from τ onwards, with the threshold for significance increased with a Bonferroni correction depending on the number of times τ had been rejected. Again, the Hamed-Kao test was used to evaluate the trend for this new guess of τ. This process was repeated until the Hamed-Kao test and Sen’s slope indicated that there was no significant movement, or the segmented *S* had become too small. In the case of the latter, the maximum number of laps was returned.

To assess whether the output was specific to the RQA pipeline the output was validated with a model-free approach using the trajectory of the manifold. A divergence metric quantified each post-switch lap’s distance from an initial centroid. The centroid was defined as the average state in the embedding of the first 15% of laps. The divergence was overlaid with the identified stability onset to check whether a plateau in divergence distance coincided with the onset of stability.

### Behavioral stability

The velocity of the mouse and its licking were originally stored as a one-dimensional list of samples. These recordings were then transformed into a structure like the representational rate maps, a matrix of shape (*L, B*), where L is the number of laps and *B* is the number of bins. For licking, an entry in this matrix was the number of licks that occurred in that spatial bin on that specific lap. For velocity, this entry was the average velocity in that bin on that lap. These two matrices were combined into a single behavioural matrix, intended to reflect how behaviour is globally modulated by different experimental conditions. The two modalities were Z-scored across rows separately before concatenating. This ensured each behavioural modality contributes equally per lap. This behavioural matrix was then transformed into a Pearson correlation matrix, that describes how similar the rodent’s behaviour was from one lap to another. This matrix was provided as an input into the same pipeline used for identifying stability in the neural representation. The output, τ, was the point at which behaviour had stabilised.

### Statistical analysis of representational and behavioural stability

The representational and behavioural stability points were grouped by experiment. The samples belonging to the expected uncertainty, unexpected uncertainty, and uncertainty interaction groups were analysed with a three-way repeated measures ANOVA, and a post-hoc paired t-test, using a Bonferroni correction for multiple comparisons. The stability identification pipeline was validated with a Monte Carlo permutation test. The representational rate maps were shuffled along the temporal axis, creating 1000 shuffled variants for each mouse in each experiment. These shuffled variants were supplied to the stability identification pipeline and their results were noted. These results were subject to the same statistical analysis described above, creating a null distribution of test statistics from the ANOVA and post-hoc test. The true T-statistics and F-statistics were compared to the null distribution of T-statistics and F-statistics. The p-value was estimated by the proportion of simulated permutations that had a test statistic greater than, or equal to, the observed data. Only the pairs of experiments that had significant relationships were subject to this comparison. The returned p-value was equal to 0.001 for all pairs, meaning that there were no instances of a more extreme test statistic in the null distribution.

## Data and code availability

Data and code are available on request.

## Acknowledgements

This work was supported by a Biotechnology and Biological Sciences Research Council-funded South West Biosciences Doctoral Training Partnership [BB/T008741/1] studentship to DG and grants to JRM (BBSRC BB/V001728/1 and MRC MR/X010910/1). Pump priming funding was received from the Alzheimer’s Research UK South West Network Centre (ARUK-NC2022-SW).

We are grateful for technical assistance from Feng Xuan, Matt Udakis, Marie Sabec, Laura Alberio and Elsa Oakes and for discussions with Peter Dayan, Matt Jones, Zach Mainen, Tony Pickering, Mark Walton and all members of Mellor group on development of the URTask.

## Contributions

### CREDIT taxonomy

Conceptualization: DG, JRM, JW; Methodology: DG, JBI, DD, JRM, JW; Software: DG, AB, CT, JW; Formal analysis: DG, AB; Investigation: DG; Resources: GL, YL; Data curation: DG; Writing – Original Draft: DG, AB, JRM, JW; Writing – Review & Editing: DG, AB, CT, JBI, YL, DD, MR, JTB, JRM, JW; Visualisation: DG, AB; Supervision: MR, JTB, JRM, JW; Funding acquisition: JRM, JW.

## Supplementary information and figures

**Table S1.** Statistical comparisons of GRAB-ACh4l fluorescence (ΔF/F) across uncertainty manipulations. GGp: Greenhouse-Geisser corrected p-value.

| Novel environment GRAB-ACh4I $\Delta F/F$ (related to Fig. 2F) | | | | | | |
| --- | --- | --- | --- | --- | --- | --- |
| RM-ANOVA | N | F | $df_{\text{Factor}}$ | $df_{\text{Error}}$ | GGp | Parital $\eta^2$ |
|  | 12 | 10.47 | 2 | 22 | 0.003 | 0.49 |
| Post-hoc tests | P | Cohen's D |  |  |  |  |
| Baseline vs. Early Post-switch | < 0.001 | 2.08 |  |  |  |  |
| Baseline vs. Late Post-switch | 0.019 | 0.79 |  |  |  |  |
| Expected uncertainty GRAB-ACh4I $\Delta F/F$ (related to Fig. 3E) | | | | | | |
| RM-ANOVA | N | F | $df_{\text{Factor}}$ | $df_{\text{Error}}$ | GGp | Parital $\eta^2$ |
|  | 7 | 28.17 | 2 | 12 | < 0.001 | 0.82 |
| Post-hoc tests | P | Cohen's D |  |  |  |  |
| Baseline vs. Early Post-switch | 0.392 | 0.35 |  |  |  |  |
| Baseline vs. Late Post-switch | 0.004 | 2.00 |  |  |  |  |
| Unexpected uncertainty GRAB-ACh4I $\Delta F/F$ (related to Fig. 4E) | | | | | | |
| RM-ANOVA | N | F | $df_{\text{Factor}}$ | $df_{\text{Error}}$ | GGp | Parital $\eta^2$ |
|  | 7 | 1.41 | 2 | 12 | 0.281 | 0.19 |
| Post-hoc tests | P | Cohen's D |  |  |  |  |
| Baseline vs. Early Post-switch | 0.178 | 0.77 |  |  |  |  |
| Baseline vs. Late Post-switch | 0.535 | 0.25 |  |  |  |  |
| Uncertainty interaction GRAB-ACh4I $\Delta F/F$ (related to Fig. 5E) | | | | | | |
| RM-ANOVA | N | F | $df_{\text{Factor}}$ | $df_{\text{Error}}$ | GGp | Parital $\eta^2$ |
|  | 8 | 4.78 | 2 | 14 | 0.058 | 0.41 |
| Post-hoc tests | P | Cohen's D |  |  |  |  |
| Baseline vs. Early Post-switch | 0.05 | 1.00 |  |  |  |  |
| Baseline vs. Late Post-switch | 0.053 | 0.82 |  |  |  |  |

**Table S2.** Statistical comparisons of pupil area (z-scored) across uncertainty manipulations. GGp: Greenhouse-Geisser corrected p-value.

| Expected uncertainty pupil area (related to Fig. 3E) |  |  |  |  |  |  |
| --- | --- | --- | --- | --- | --- | --- |
| RM-ANOVA | N | F | *df*<sub>Factor</sub> | *df*<sub>Error</sub> | GGp | Parital $\eta^2$ |
|  | 7 | 31.58 | 2 | 12 | 0.001 | 0.84 |
| Post-hoc tests | P | Cohen's D |  |  |  |  |
| Baseline vs. Early Post-switch | 0.052 | 0.92 |  |  |  |  |
| Baseline vs. Late Post-switch | 0.003 | 2.09 |  |  |  |  |
| Unexpected uncertainty pupil area (related to Fig. 4E) |  |  |  |  |  |  |
| RM-ANOVA | N | F | *df*<sub>Factor</sub> | *df*<sub>Error</sub> | GGp | Parital $\eta^2$ |
|  | 7 | 4.56 | 2 | 12 | 0.046 | 0.43 |
| Post-hoc tests | P | Cohen's D |  |  |  |  |
| Baseline vs. Early Post-switch | 0.039 | 1.19 |  |  |  |  |
| Baseline vs. Late Post-switch | 0.116 | 0.69 |  |  |  |  |
| Uncertainty interaction pupil area (related to Fig. 5E) |  |  |  |  |  |  |
| RM-ANOVA | N | F | *df*<sub>Factor</sub> | *df*<sub>Error</sub> | GGp | Parital $\eta^2$ |
|  | 8 | 6.92 | 2 | 14 | 0.018 | 0.50 |
| Post-hoc tests | P | Cohen's D |  |  |  |  |
| Baseline vs. Early Post-switch | 0.421 | 0.30 |  |  |  |  |
| Baseline vs. Late Post-switch | 0.004 | 1.72 |  |  |  |  |

**Table S3.** Statistical comparisons of manifold tortuosity across uncertainty manipulations. EU: Expected Uncertainty; UU: Unexpected Uncertainty; UI: Uncertainty Interaction.

| Neural Manifold Tortuosity |  |  |  |  |  |  |
| --- | --- | --- | --- | --- | --- | --- |
| RM-ANOVA | N | F | $df_{\text{Factor}}$ | $df_{\text{Error}}$ | P | Parital $\eta^2$ |
|  | 4 | 8.72 | 2 | 6 | 0.016 | 0.63 |
| Post-hoc tests | P | Cohen's D |  |  |  |  |
| EU vs UU | 0.203 | -1.34 |  |  |  |  |
| UI vs UU | 0.424 | 1.57 |  |  |  |  |
| UI vs EU | 0.127 | 2.77 |  |  |  |  |

**Table S4.** Statistical comparisons of stabilisation time across uncertainty manipulations. EU: Expected Uncertainty; UU: Unexpected Uncertainty; UI: Uncertainty Interaction.

| Neural stability |  |  |  |  |  |  |
| --- | --- | --- | --- | --- | --- | --- |
| RM-ANOVA | N | F | *df*<sub>Factor</sub> | *df*<sub>Error</sub> | P | Parital $\eta^2$ |
|  | 4 | 65.75 | 2 | 6 | <0.001 | 0.79 |
| Post-hoc tests | P | Cohen's D |  |  |  |  |
| EU vs UU | 0.005 | 2.78 |  |  |  |  |
| UI vs UU | 0.004 | 5.12 |  |  |  |  |
| UI vs EU | 0.670 | 0.64 |  |  |  |  |
| Behavioral stability |  |  |  |  |  |  |
| RM-ANOVA | N | F | *df*<sub>Factor</sub> | *df*<sub>Error</sub> | P | Parital $\eta^2$ |
|  | 4 | 20.06 | 2 | 6 | 0.002 | 0.67 |
| Post-hoc tests | P | Cohen's D |  |  |  |  |
| EU vs UU | 0.027 | 2.43 |  |  |  |  |
| UI vs UU | 0.017 | 3.59 |  |  |  |  |
| UI vs EU | 1.000 | 0.09 |  |  |  |  |

**Figure S1.**
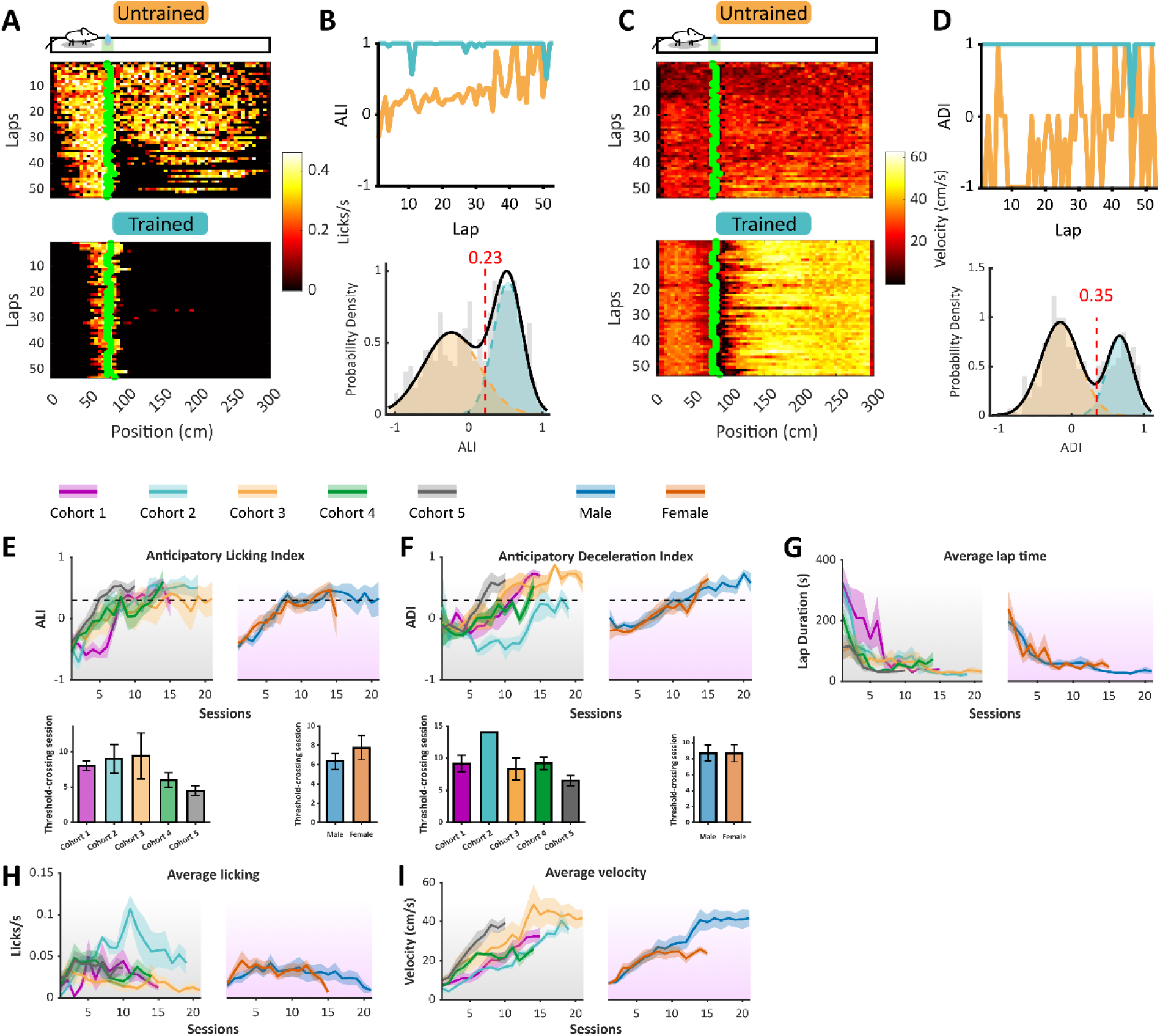
Consistent task learning across sex and cohort of mice. (A) Representative heatmaps of licking behaviour across laps in an early (untrained) and late (trained) session from a single mouse. Each row represents a lap, with colour indicating licking rate. Green dots mark the reward location. Schematic above shows track layout and reward location in green. (B) Top: Anticipatory licking index (ALI) for untrained (orange) and trained (blue) sessions in (A), showing increased licking near the reward site after training. Bottom: Gaussian mixture model (GMM) of licking indices reveals two distinct behavioural populations; the red dashed line indicates the classification threshold separating untrained and trained distributions. (C) Velocity heatmaps from the same sessions and mouse as in (A), showing approach behaviour to the reward (green dots) across laps. (D) Top: Anticipatory deceleration index (ADI) for untrained (orange) and trained (blue) sessions in (C), showing enhanced slowing near the reward with training. Bottom: GMM analysis identifies two distributions; the red dashed line denotes the intersection used to define training threshold. (E) Top left: ALI across sessions in all cohorts (grey background), with a 0.23 dashed line denoting the threshold for trained behaviour. Top right: Same analysis split by sex (purple background). Bottom: Bar graphs showing the average session in which each cohort (left) or sex (right) reached the threshold; no significant differences observed. (F) Same as in (E), but for the ADI. Threshold set to 0.35 indicated in D.; no significant differences across cohorts or sex in threshold crossing sessions (repeated measures ANOVA with Bonferroni-corrected post hoc t-tests, N = 27). (G–I) Average licking rate (G), average velocity (H), and average lap time (I) across training sessions, split by cohort (left, grey background) and by sex (right, purple background). No consistent differences observed across groups (repeated measures ANOVA with Bonferroni-corrected post hoc t-tests, N = 27).

**Figure S2.**
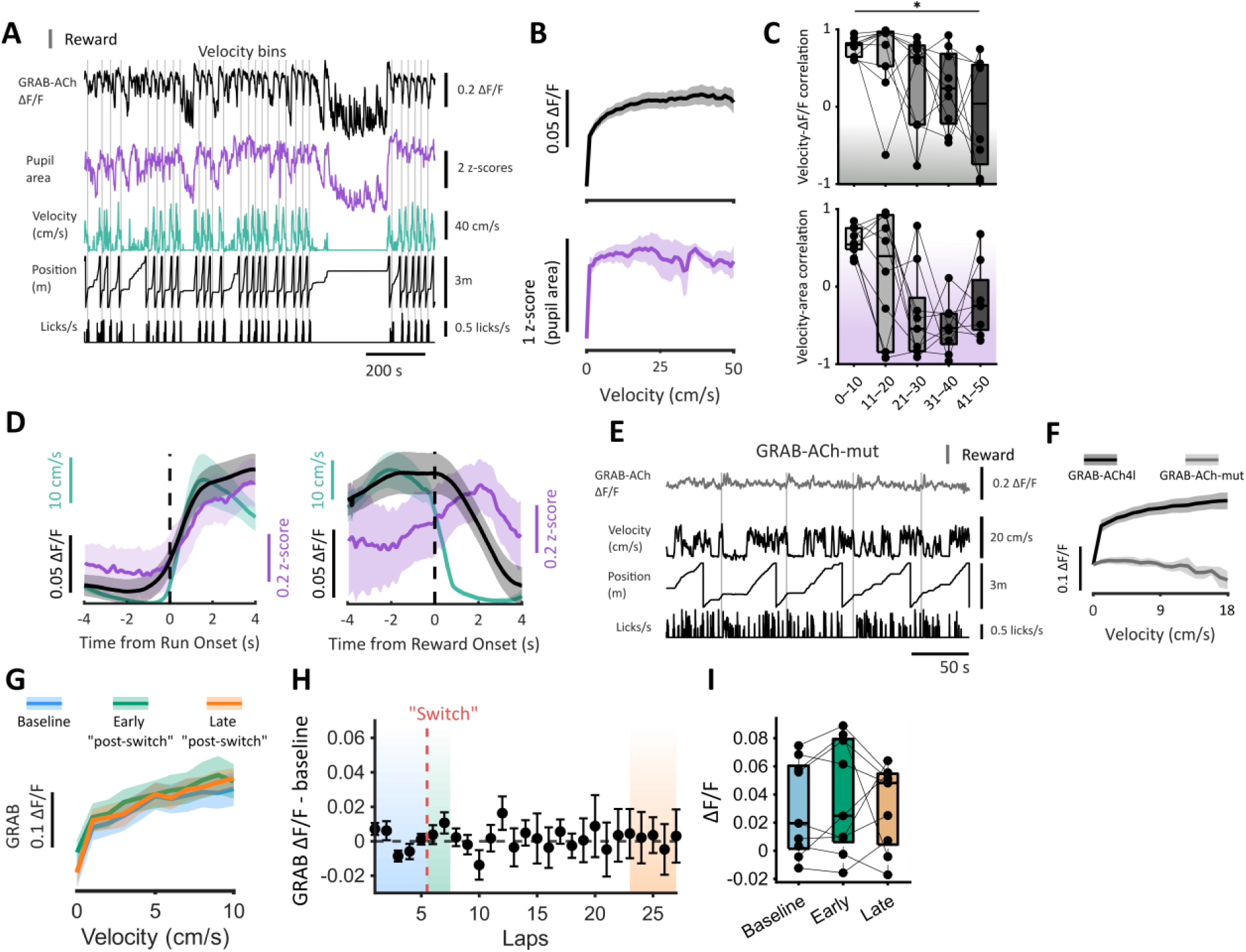
Characterisation of GRAB-ACh sensor. (A) GRAB-ACh4l ΔF/F, pupil area, velocity, track position, and lick traces for an example recording session; reward deliveries marked by vertical grey lines. (B) Velocity-binned GRAB-ACh4l ΔF/F in dRSC layer II/III. (C) Correlations between velocity and ΔF/F (top) or pupil area (bottom) across velocity bins (0–50 cm/s). Green and blue shading indicate GRAB-ACh4l and pupil data, respectively. Repeated measures ANOVA with Bonferroni-corrected post hoc comparisons. (D) (Left) average peri-run onset traces (±4 s) for ΔF/F (green), velocity (pink), and pupil area (blue). (Right) average peri-reward onset traces (±4 s) for ΔF/F (green), velocity (pink), and pupil area (blue). (E) Example traces from mouse expressing non-responsive GRAB-ACh-mut sensor. (F) ΔF/F across velocity bins comparing GRAB-ACh4l (green) and GRAB-ACh-mut (grey). (G) ÄF/F binned by velocity during a control session (no switch). Separate lines indicate baseline (blue), early post-switch (green), and late post-switch (orange) epochs. “Switch” time is selected as the 6^th^ imaged lap per session for each mouse. N = 11. (H) Lap-by-lap baseline-subtracted ΔF/F between 0-10cm/s during a control session. Red dashed line indicates 6^th^ trial for each mouse. Shaded regions denote baseline (blue), early “post-switch” (green), and late “post-switch” (orange) epochs. Dashed lines indicate zero. (I) Mean ΔF/F across behavioural epochs (baseline, early, and late). Repeated-measures ANOVA with Greenhouse–Geisser correction was used to assess main effects of epoch, followed by Dunnett-adjusted post hoc comparisons testing early and late epochs against baseline.

**Figure S3.**
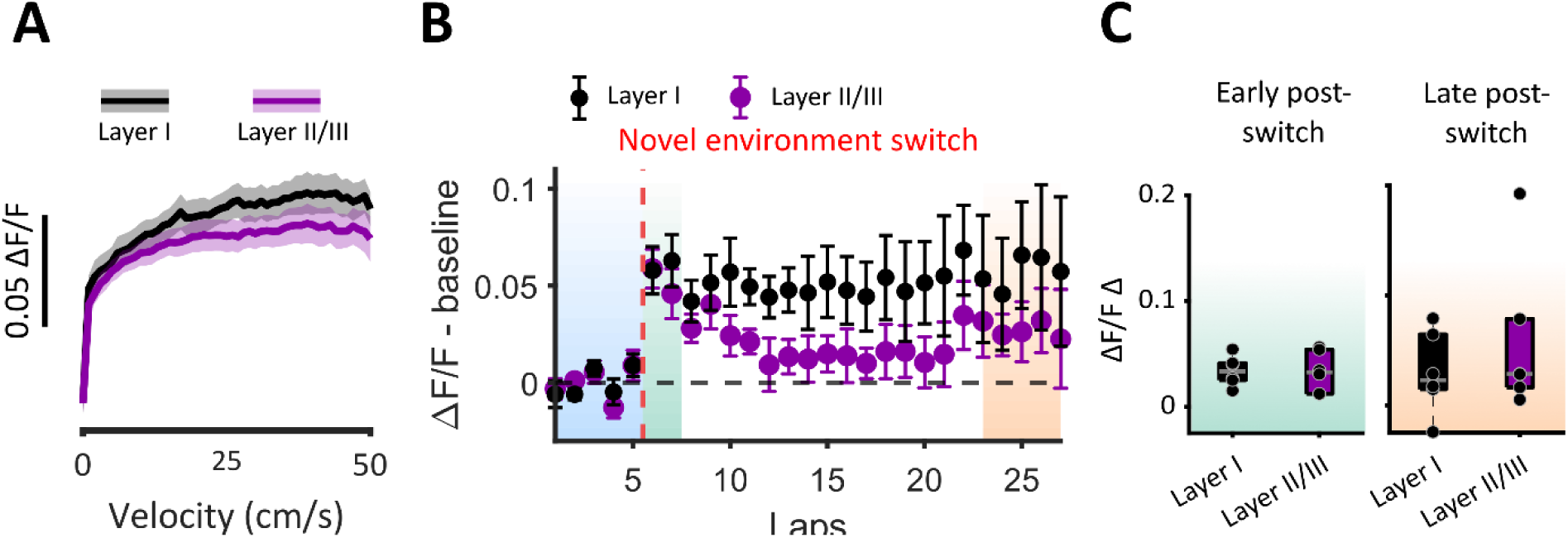
No layer dependency of acetylcholine release in dRSC. (A) Velocity-binned GRAB-ACh4l ΔF/F layers I vs. II/III. (B) Lap-by-lap time course of baseline-subtracted ΔF/F aligned to the novel environment switch. Shaded areas indicate the baseline (blue), early post-switch (light green), and late post-switch (orange) analysis windows. Red dashed line indicates switch to novel environment. Dashed line indicates 0. (C) Change in ΔF/F from baseline in early (light green shaded) and late post-switch (orange shaded) periods, comparing layer I and layer II/III for novel environment switch.

**Figure S4.**
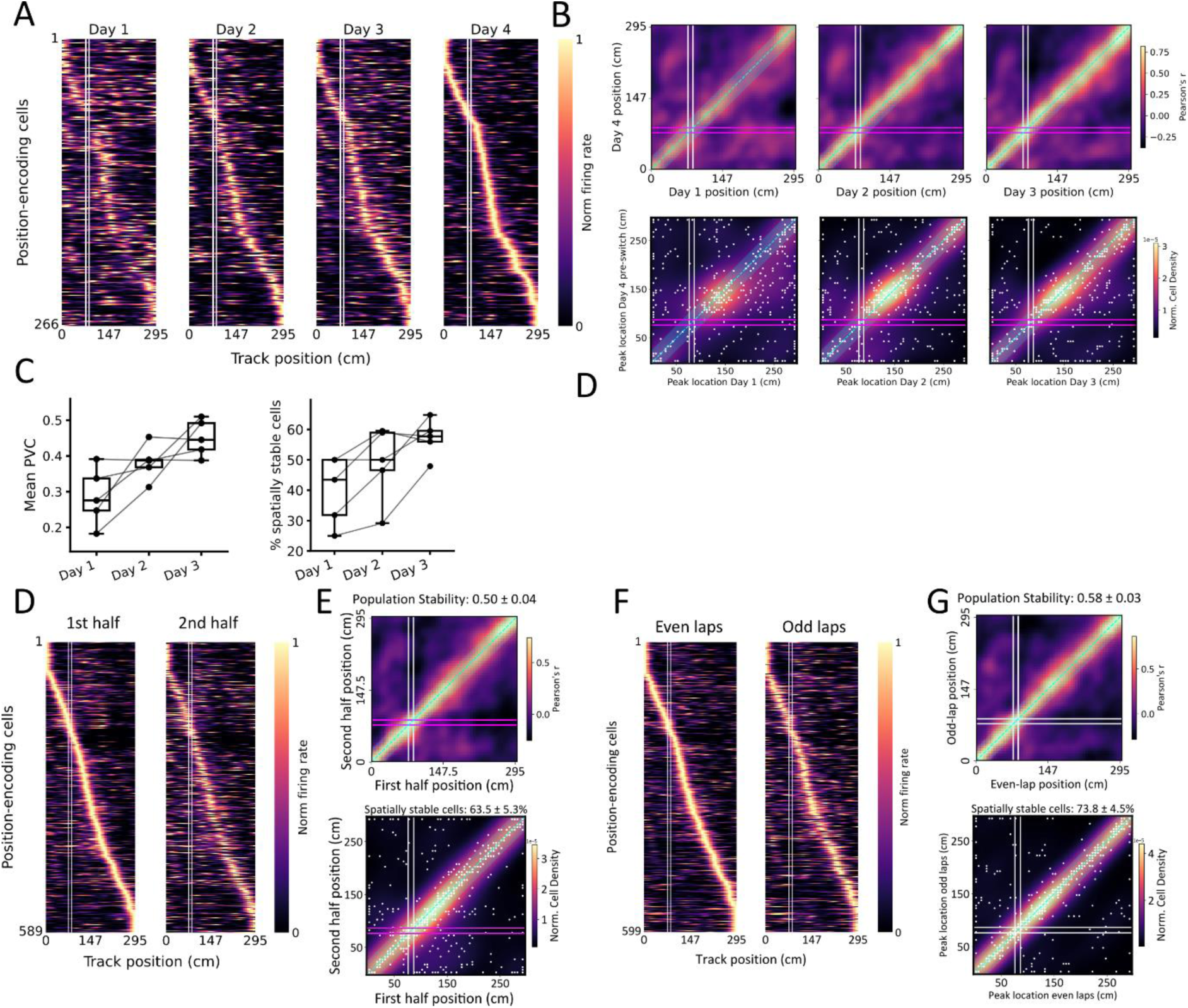
Stability of spatial representations in dRSC. (A) Normalized activity maps of example position-encoding cells across training, ordered by peak activity on the final day. White lines indicate reward location. (B) (Top) population vector correlations (PVCs) between final session (day 4) and earlier sessions. Reward locations marked (white lines). Blue shading around the unity line indicates ±25 cm tolerance for spatial stability. (Bottom) Galaxy plots indicating cell-level stability between final session and earlier sessions. Scatter points represent peak positions, overlaid on a kernel density estimate (KDE) heat map. Cells classified as spatially stable if in cyan diagonal area (peak positions within ±25 cm across sessions). (C) Quantification of PVCs across sessions (left) and percentage of spatially stable cells (right). (D) Within-session stability in well-trained mice. Left: activity maps from the first half of laps within a recording session, ordered by peak position. Right: activity maps from the second half, ordered by first-half peaks. The reward location is marked by white lines. (E) Top: PVCs comparing the first versus the second half of laps. Bottom: galaxy plot showing peak activity locations for all cells. Scatter points represent peak positions, overlaid on a kernel density estimate (KDE) heat map. Cells classified as spatially stable (peak positions within ±25 cm of unity between halves, depicted as a cyan dashed line and shading) are highlighted in green. (F) Odd versus even lap comparison of normalized activity maps. (G) PVCs (top) and galaxy plot (bottom) for odd versus even laps, as in (E).

**Figure S5.**
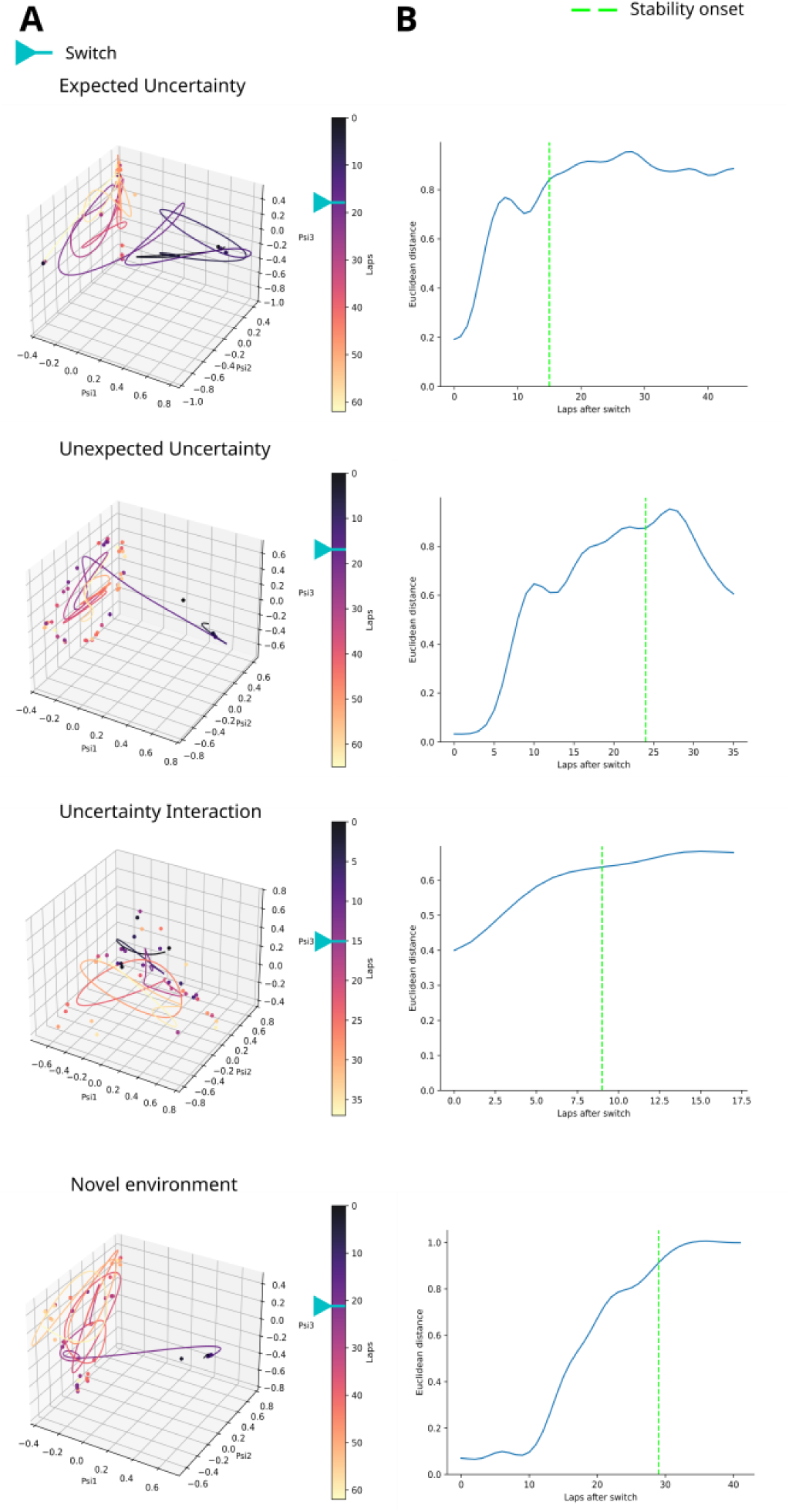
Behavioural manifolds and model-free validation of stability analysis. (A) Example diffusion maps of behaviour across switch conditions. (B) The Euclidean distance travelled by the manifold from an initial post-switch centroid.

